# Iron-sequestering nanocompartments as multiplexed Electron Microscopy gene reporters

**DOI:** 10.1101/516955

**Authors:** Felix Sigmund, Susanne Pettinger, Massimo Kube, Fabian Schneider, Martina Schifferer, Michaela Aichler, Steffen Schneider, Axel Walch, Thomas Misgeld, Hendrik Dietz, Gil G. Westmeyer

## Abstract

Multi-colored gene reporters such as fluorescent proteins are indispensable for biomedical research, but equivalent tools for electron microscopy (EM), a gold standard for deciphering mechanistic details of cellular processes^1,2^ and uncovering the network architecture of cell-circuits^3,4^, are still sparse and not easily multiplexable. Semi-genetic EM reporters are based on the precipitation of exogenous chemicals^5–9^ which may limit spatial precision and tissue penetration and can affect ultrastructure due to fixation and permeabilization. The latter technical constraints also affect EM immunolabeling techniques^10–13^ which may furthermore be complicated by limited epitope accessibility. The fully genetic iron storage protein ferritin generates contrast via its electron-dense iron core^14–16^, but its small size complicates differentiation of individual ferritin particles from cellular structures. To enable multiplexed gene reporter imaging via conventional transmission electron microscopy (TEM), we here introduce the encapsulin system of *Quasibacillus thermotolerans* (Qt) as a fully genetic iron-biomineralizing nanocompartment. We reveal by cryo-electron reconstructions that the Qt monomers (QtEnc) self-assemble to nanospheres with T=4 icosahedral symmetry and an ~44 nm diameter harboring two putative pore regions at the fivefold and threefold axes. We furthermore show that the native cargo (QtIMEF) auto-targets to the inner surface of QtEnc and exhibits ferroxidase activity leading to efficient iron sequestration inside mammalian cells. We then demonstrate that QtEnc can be robustly differentiated from the non-intermixing encapsulin of *Myxococcus xanthus*^17^ (Mx, ~32 nm) via a deep-learning model, thus enabling automated multiplexed EM gene reporter imaging in mammalian cells.

---

Encapsulins are a class of proteinaceous spherical nanocompartments naturally occurring in bacteria and archaea, so far described as icosahedral structures with either T=1 (60 subunits, ~18 nm diameter) or T=3 (180 subunits, ~30 nm) symmetry, which can encapsulate cargo proteins with a wide range of functions^18–21^. It has also been shown that foreign cargos such as fluorescent proteins or enzymes can be genetically targeted into the encapsulin lumen in bacterial and mammalian hosts^22–26^.

## Expression, assembly, and cargo-loading of QtEnc in mammalian cells

To express the encapsulin genes from Qt in comparison to Mx in mammalian cells, we used mammalian expression constructs for the shell proteins (QtEnc, MxEnc) and the native cargos (QtIMEF, MxBCD as a P2A construct) (**Fig. 1a,b; Supplementary Table 1**). Blue Native PAGE (BN-PAGE) analysis of lysates from HEK293T cells co-expressing MxEnc+MxBCD showed two bands corresponding to the T=1 and T=3 assemblies, as we have previously shown^26^. In contrast, expression of QtEnc+QtIMEF yielded a single band running slightly higher than the band corresponding to the T=3 assembly of MxEnc (**Fig. 1c**). To further investigate the size differences between QtEnc and MxEnc, we purified QtEnc nanoshells from HEK293T cells via their external FLAG-epitope and performed dynamic light scattering (DLS) yielding diameters of 39 ± 14 nm for QtEnc and 40 ± 2 nm for QtEnc+QtIMEF (**Supplementary Fig. 1**).

**Fig. 1.**
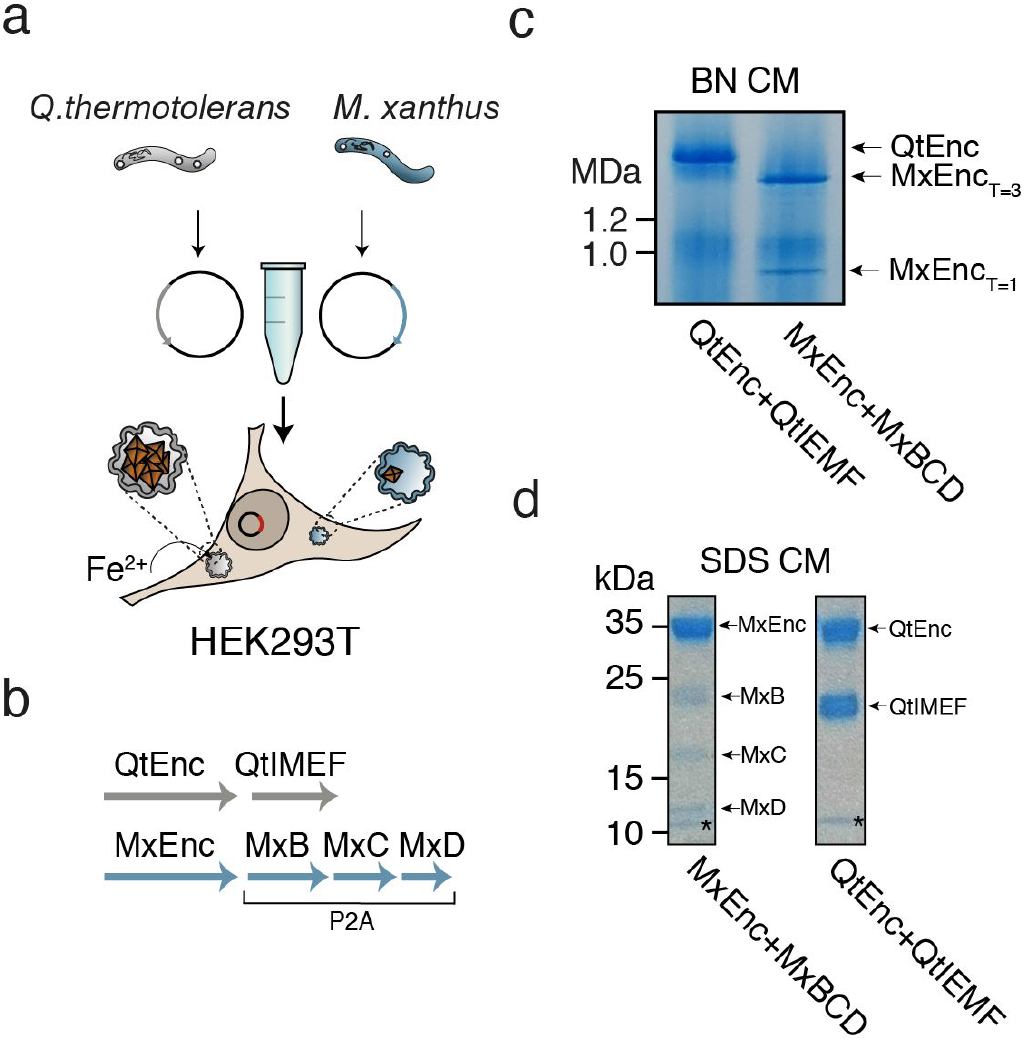
Encapsulins of different bacterial species express, assemble and load cargo proteins in mammalian cells. **a**, Schematic depiction of the heterologous expression and self-assembly of encapsulins in mammalian cells. **b**, Scheme of the mammalian expression constructs encoding the shell and cargo proteins of *Quasibacillus thermotolerans* (QtEnc+QtIMEF) and *Myxococcus xanthus* (MxEnc+MxBCD). Please see Supp. Table 1 for details. **c**, Coomassie-stained Blue Native PAGE (BN CM) gel loaded with cell lysates of HEK293T cells expressing QtEnc+QtIMEF and MxEnc+MxBCD. **d**, Coomassie-stained SDS-PAGE (SDS CM) gel of both encapsulin:cargo systems purified via their FLAG epitope showing bands for the respective shell and co-precipitated cargo proteins. The dye running front is visible above the 10 kDa marker band on both lanes (*).

To confirm that auto-targeting of QtIMEF to QtEnc still occurs in mammalian cells, we purified encapsulins from HEK293T cells via their external FLAG-epitope and evaluated their protein contents via SDS-PAGE. We detected two bands corresponding to the shell protein and the cargo (QtEnc: 32 kDa, QtIMEF: 23 kDa)^20^ (**Fig. 1d**). We then used densitometric SDS gel quantification to determine the ratio of shell to cargo for both encapsulin systems, and estimated that the MxEnc shell accounted for 47 ± 4 % of total protein and the three cargo proteins for 53 ± 4 % (MxB: 86 ± 3 molecules; MxC: 93 ± 9 molecules; MxD: 50 ± 15 molecules; *n*=3), whereas for QtEnc 51 ± 1% of total protein belonged to the shell and 49 ± 1 % (231 ± 5 molecules; *n*=3) to the QtIMEF cargo (**Fig. 1d, Supplementary Table 2**).

## Structural characterization of QtEnc without and with QtlMEF cargo

To gain insights into the structural characteristics of Qt encapsulin, we conducted single particle cryo-EM analysis on purified QtEnc nanocompartments sans and with encapsulated QtIMEF cargo (**Fig. 2, Supplementary Fig. 2,3**). Analysis of QtEnc without cargo revealed hollow compartments with T=4 icosahedral symmetry consisting of 240 subunits (**Fig. 2a, Supplementary Fig. 2a-f**). We found a circular electron-sparse region with a diameter of about 1 nm on the five-fold symmetry axis, a triangular shaped region at the threefold axis, and an additional electron-sparse cleft on the twofold symmetry center (**Fig.2a and Supplementary Fig. 2g-i**). Further structural analysis performed on QtEnc particles loaded with QtIMEF showed a maximum outer diameter of ~44 nm and additional electron densities in ~5 nm distance to the inner shell surface, corresponding to the docked cargo (**Fig. 2b; Supplementary Fig. 3**). This observation is in agreement with our earlier finding that cargo auto-targeting still occurs efficiently upon heterologous expression in mammalian cells (**Fig. 1d**).

**Fig. 2.**
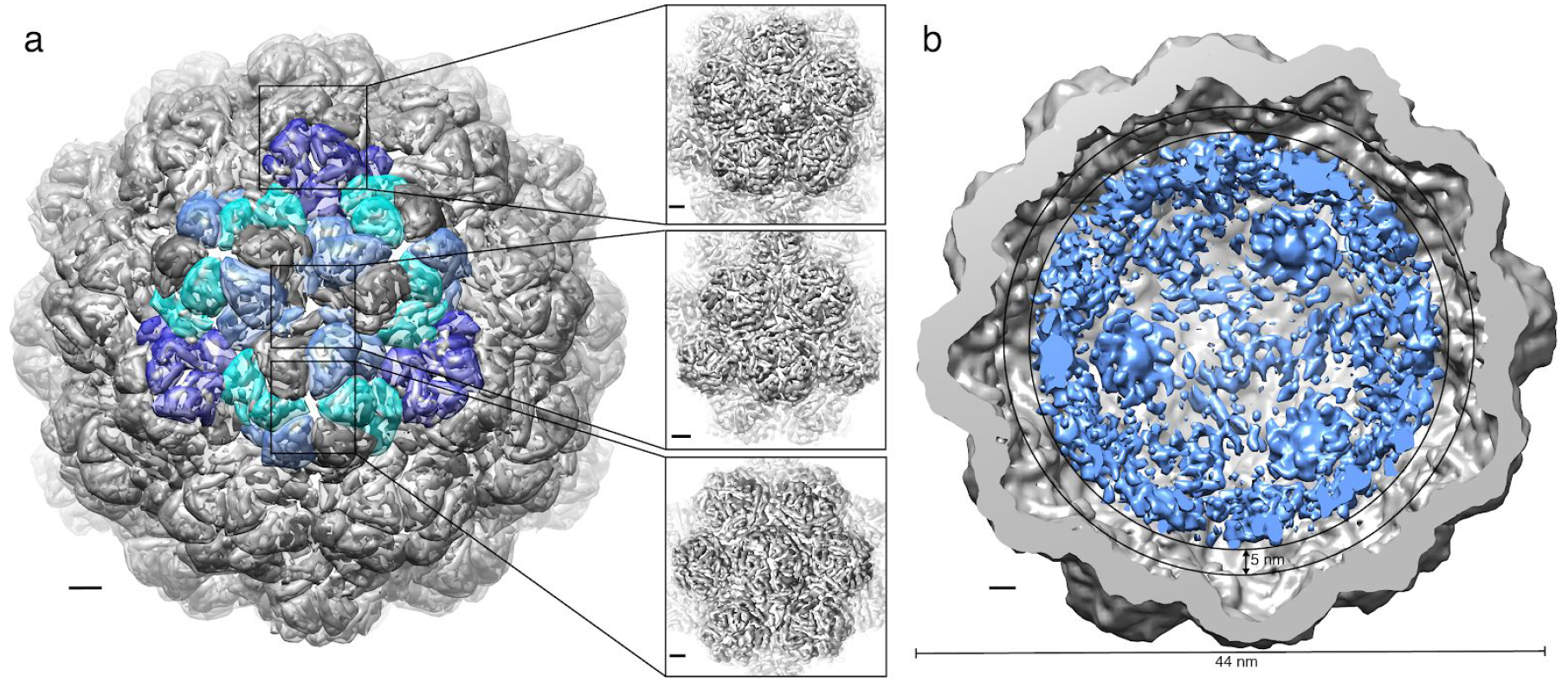
Cryo-EM reconstruction of QtEnc reveals icosahedral T=4 symmetry. **a**, Segmented electron density map of QtEnc without cargo purified from mammalian cells. The four different monomer conformations are colored according to their interconnectivity and position. The fivefold centers are on opposite sides of a threefold center with two monomers in between, which indicates a T=4 icosahedral symmetry of the shell. Boxes show zoomed-in views of the fivefold, threefold, and twofold symmetry centers. The resolution of the map is 7.6 Å; scale bars represent 2 nm. **b**, Cutaway view through the maximum diameter of QtEnc (44 nm) showing the shell (grey) and co-expressed QtIMEF cargo (blue) at different electron densities. A gap of 5 nm is apparent between cargo and shell. The resolution of the map is 11 Å.

## Iron biomineralization inside QtEnc+QtIMEF expressed in mammalian cells

Since iron accumulation can substantially enhance contrast in TEM^16^, we next evaluated how efficiently QtIMEF biomineralizes iron inside QtEnc within mammalian cells as compared to the ferroxidase-driven iron-mineralization inside MxEnc. We thus expressed QtEnc+QtIMEF or MxEnc+MxBCD, supplemented different concentrations of ferrous ammonium sulfate (FAS) for 36 hours, and investigated iron loading via 3,3’-Diaminobenzidine-enhanced Prussian Blue (DAB PB) stained BN-PAGE of cell lysates. We detected only weak light blue bands for MxEnc+MxBCD for all tested FAS concentrations (due to residual Coomassie dye in the cathode buffer), while QtEnc+QtIMEF showed distinct brown precipitates already for the lowest FAS supplement of 0.125 mM (**Fig. 3a**).

**Fig. 3.**
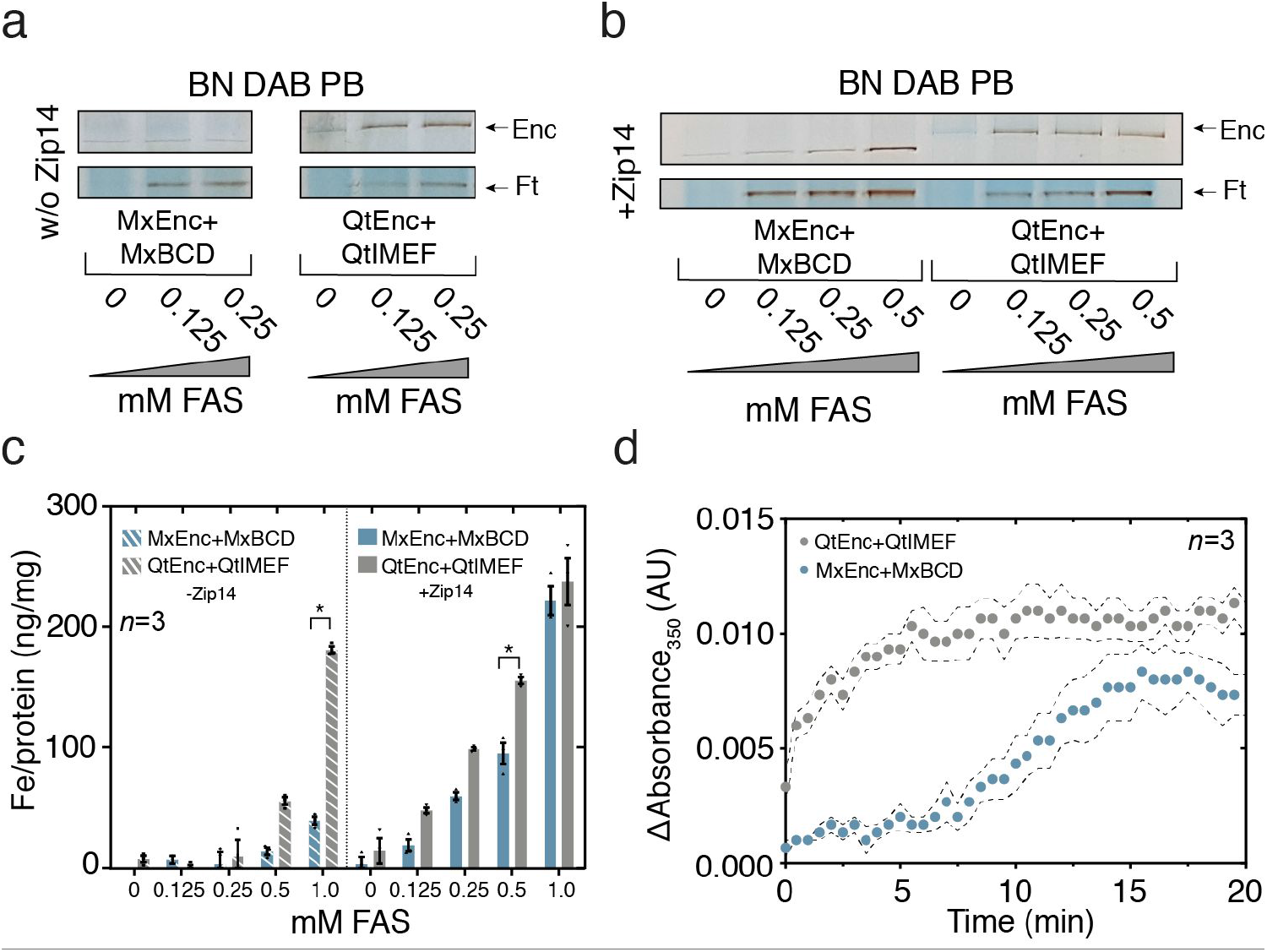
QtEnc+QtIMEF biomineralizes iron more efficiently than MxEnc+MxBCD. **a,b**, DAB-enhanced Prussian Blue-stained BN-PAGE (BN DAB PB) analysis of whole-cell lysates. HEK293T cells expressing MxEnc+MxBCD or QtEnc+QtIMEF were supplemented with different concentrations of ferrous ammonium sulfate (FAS) for 36 h with or without co-transfection of the iron-transporter Zip14. The upper panels show bands corresponding to the assembled encapsulin constructs, whereas the lower panels correspond to endogenous ferritin (Ft). Brown precipitates indicate iron-loaded protein complexes. In panel a, the DAB-enhancement reaction was stopped after 2 h, whereas in b the reaction was stopped after 30 min. **c**, Total iron contents normalized to total protein concentration (OD_280_) in cell lysates of HEK293T cells (3 independent biological replicates) expressing QtEnc+QtIMEF or MxEnc+MxBCD with or without Zip14 after supplementation of different FAS concentrations for 36 h. Stars indicate *p* values < 0.05 in post hoc tests of a 2-way ANOVA (see Supp. Table 3 for all results of the statistical tests). **d**, Spectroscopic monitoring of iron biomineralization inside QtEnc+QtIMEF and MxEnc+MxBCD. 50 Fe^2+^ ions per cargo monomer were added to 16 μM of cargo monomer (QtIMEF or MxBCD) inside the respective encapsulin shells and absorbance was recorded at 350 nm for 20 min at room temperature after a deadtime due to pipetting and mixing of ~1 min. Absorbance traces were corrected for iron auto-oxidation by subtracting the absorbance trace of Fe^2+^ in solution. Traces represent the mean ± SEM.

When we co-expressed the iron importer Zip14 to increase cellular iron uptake^26,27^, we detected iron biomineralization within MxEnc+MxBCD for 0.5 mM FAS as we have shown previously^26^, whereas iron-loading of QtEnc+QtIMEF was already saturated at 0.125 mM (**Fig. 3b**). The BN-PAGE analysis also revealed strong iron loading of native ferritin in cells co-expressing MxEnc+MxBCD, whereas cells expressing QtEnc+QtIMEF showed weaker ferritin bands in comparison (**Fig. 3b**, lower panel). This result indicates that QtEnc competes more effectively with endogenous ferritin for iron than MxEnc. To assess possible effects of FAS supplementation on cell viability, we conducted a sensitive luciferase-based assay that showed no significant decrease in viability upon treatment with up to 0.5 mM FAS for all tested genetic constructs (**Supplementary Fig. 4** and **Supplementary Table 3** for complete statistical analysis).

To quantify the influence of QtEnc+QtIMEF or MxEnc+MxBCD overexpression on the whole cell iron content, we conducted a colorimetric iron assay on cell lysates. Without co-expression of Zip14, we observed significant effects for encapsulin type (2-way ANOVA, *p*<0.0001), and FAS concentration *p*<0.0001), as well as a significant interaction (*p*<0.0001) (**Fig. 3c**). At 0.5 mM FAS, cells overexpressing QtEnc+QtIMEF but not MxEnc+MxBCD showed significantly increased iron accumulation compared to cells without FAS supplementation (post hoc test with Bonferroni correction, *p*=0.0166 and *p*=0.5833, *n*=3). In addition, QtEnc+QtIMEF expression led to a significantly higher total iron content than expression of MxEnc+MxBCD at 1 mM FAS (*p*<0.0001, please also refer to **Supplementary Table 3**). For cells co-expressing Zip14 and the shell/cargo constructs, we also observed significant effects for the type of encapsulin (2-way ANOVA, *p*<0.0001) and FAS concentration (*p*=0.0001), but no significant interaction between the two factors (*p*=0.0775) (**Fig. 3c**). The increase in total iron content compared to untreated cells was significant for expression of either MxEnc+MxBCD or QtEnc+QtIMEF at FAS concentrations of 0.25 mM and higher (post hoc test with Bonferroni correction, *p*<0.05 for 0.25, 0.5, and 1 mM FAS). We also observed a significantly higher iron accumulation upon expression of QtEnc+QtIMEF compared to MxEnc+MxBCD at a FAS concentration of 0.5 mM (*p*=0.0048), whereas there was no significant difference between the two encapsulin systems at all other tested FAS concentrations (please refer to **Supplementary Table 3** for a list of all *p*-values).

To further quantify the number of iron atoms per encapsulin assembly, we conducted ICP-MS analysis for QtEnc+QtIMEF and MxEnc+MxBCD purified from HEK293T cells co-expressing low-level Zip14. We found that, on average, QtEnc+QtIMEF contained ~35000 iron atoms per shell, whereas MxEnc+MxBCD only contains ~19000 Fe (**Supplementary Table 2**).

## Ferroxidase activity of encapsulin systems

To further characterize the iron biomineralization efficiency of QtEnc+QtIMEF and MxEnc+MxBCD, we purified the two encapsulin shells from medium-scale batches of HEK293T cells co-expressing the respective cargos and ran an established ferroxidase assay^28^ (**Fig. 3d**). We added 50 Fe^2+^ per cargo protein to either encapsulin system and monitored the absorbance at 350 nm for 20 min. For QtEnc+QtIMEF, we saw a fast initial increase in absorbance, which reached a plateau after ~5 min. For MxEnc+MxBCD, we repeatedly observed a delay of ~6 min before absorbance increased and a plateau was reached after ~15 min (**Fig. 3d**, *n*=3 measurements). Compared to QtEnc+QtIMEF, the ferroxidase activity of MxEnc+MxBCD showed a slower onset and led to a lower total absorbance after reaching a plateau, which suggests a faster activity of QtIMEF compared to MxBCD.

## Multiplexed gene reporter imaging in TEM

We then sought to test the multiplexing capability of the two encapsulin systems for cellular TEM. We thus expressed both encapsulin:ferroxidase systems with low levels of Zip14 and 0.5 mM FAS and compared their contrast in TEM to overexpressed ferritin, which was previously suggested as an EM-marker^14–16^. Both encapsulin:ferroxidase systems could be clearly detected as individual nanospheres dispersed throughout the cytosol and filled with electron-rich cores. In distinction, ferritin particles could not be identified within the cells on our TEM system (**Fig. 4a**). We manually measured the diameters of cytosolic QtEnc and MxEnc and found a significant size difference of the nanospheres with diameters of 38.3 ± 1.5 nm and 32.2 ± 2.0 nm respectively (two-tailed t-test, *p*<0.0001, *n*=100), showing that the two different encapsulin nanoshells could be differentiated in TEM sections based on their size (**Fig. 4b**).

**Fig. 4.**
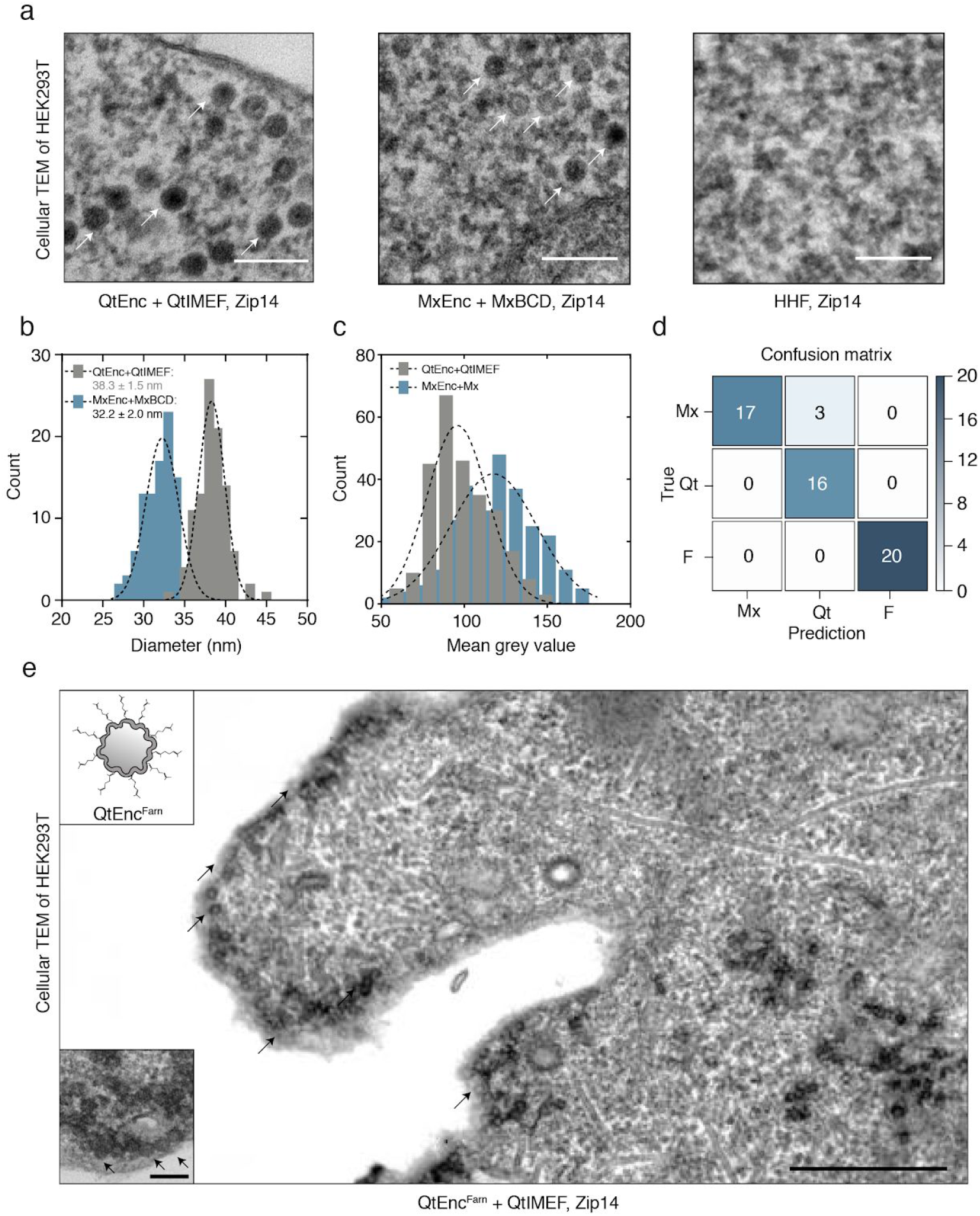
QtEnc and MxEnc serve as a multiplexed genetically encoded EM reporters. **a**, TEM images from HEK293T cells expressing either QtEnc+QtIMEF, MxEnc+MxBCD, or HHF as a control. Cells were supplemented with 0.5 mM FAS for 36 h and co-expressed low levels of Zip14. Scale bar represents 100 nm. White arrows highlight encapsulin nanoshells. **b**, Histogram of the diameters for QtEnc+QtIMEF and MxEnc+MxBCD obtained from manual segmentation of 100 nanospheres from TEM images of HEK293T cells. **c**, Histogram of mean gray values obtained from a manual segmentation of QtEnc+QtIMEF and MxEnc+MxBCD in the TEM images. **d**, Performance of the automated TEM image classification using a deep learning approach. The aggregated confusion matrix was obtained from 5-fold cross-validation run with each full-sized image treated as one sample. **e**, Full frame EM image of a HEK293T cell expressing the farnesylated Qt variant (QtEnc^Farn^) with QtIMEF cargo and co-expression of low-level Zip14. Black arrows indicate encapsulins close to the plasma membrane. Scale bar represents 500 nm. The inset at the upper left corner shows a scheme of a farnesylated QtEnc shell. Bottom left corner: Close-up of an exemplary region around the plasma membrane with black arrows pointing to the farnesylated encapsulins. Scale bar represents 100 nm.

From our earlier observations that the two encapsulin systems showed different iron mineralization behavior, we furthermore reasoned that Qt nanospheres should yield higher contrast on TEM images. Thus, we manually segmented and measured mean gray values of the two encapsulin systems and found that QtEnc nanospheres appeared significantly darker on the TEM images (two-tailed t-test, *p*<0.0001, *n*=266), which is in line with our findings that QtEnc+QtIMEF exhibited more efficient iron biomineralization and has a larger iron loading capacity compared to MxEnc+MxBCD (**Fig. 4c**).

To test whether the two encapsulin systems showed sufficiently distinct TEM features for a fully automated image classification, we adapted a pre-trained residual neural network^29^ (**Supplementary Fig. 7**) to differentiate TEM images of HEK293T cells expressing one of the three iron-storing complexes (QtEnc+QtIMEF, MxEnc+MxBCD, HHF). We compared the results in a 5-fold cross-validation scheme (**Fig. 4d**) showing that the network exceeds human performance (**Supplementary Fig. 7b**). We also expressed both encapsulin systems together and showed that nanoshells of both sizes could be clearly differentiated on TEM images even without iron supplementation. This also indicated that the shell-forming monomer subunits do not intermix with each other (**Supplementary Fig. 6c**), which was further substantiated by the distinct band patterns on BN-PAGE (**Supplementary Fig. 6d**). Furthermore, we investigated whether we could genetically modify encapsulins to target the cell membrane by adding a minimal farnesylation signal (GCMSCKCVLS, C-terminal residues 281-290 of rat H-Ras^30^) on the surface-facing C-terminus of the QtEnc monomers (QtEnc^Farn^). Upon co-expression of this variant with QtIMEF, we observed strongly contrasted encapsulins located at the plasma membrane (**Fig. 4e**).

In summary, we demonstrated that the encapsulins of Qt and Mx serve as a potent set of EM gene reporters (“EMcapsulins”) that enable multiplexed detection in mammalian cells by conventional TEM. Via cryo-EM reconstructions, we identified QtEnc as the first encapsulin with T=4 icosahedral symmetry resulting in a diameter of ~44 nm which can be robustly differentiated from the smaller T=3 Mx encapsulin (~32 nm) in TEM. The sharp size distribution of both encapsulin types highlights the superior precision in spatial localization as compared to techniques based on triggered precipitation of exogenous dyes^5–7^. In addition, the QtIMEF cargo proved to be an efficient ferroxidase even at low iron supplementation, which is notable because it does not possess the canonical iron-coordinating residues usually found in four-helix bundle proteins^20^.

In comparison to the strong contrast provided by the tens-of-nanometer-sized electron-dense cores inside the encapsulin shells, we could not detect ferritin particles despite strong overexpression and ensuring their iron-loading via coexpression of Zip14. Moreover, in distinction to bioorthogonal EMcapsulins, ferritin-based EM markers are essentially indistinguishable from endogenous ferritin whose expression is furthermore upregulated upon iron supplementation resulting in increased background signal.

We proved the robustness of contrast generation by EMcapsulins via a deep neural net-based classification system, which could reliably distinguish the two variants in cellular TEM images in a fully automated manner. Similar to the wide color spectrum of fluorescent proteins, we anticipate that the rich geometric feature space of EMcapsulins can be further expanded by adding encapsulin variants with additional sizes, different intraluminal content (*e.g*., fluorescent proteins for correlative detection or enzymes such as APEX ^26^), patterning or intracellular localization.

We expect that multiplexed imaging with EMcapsulins will be helpful for testing mechanistic hypotheses about multi-component cellular machinery and valuable for complementing EM connectome data with information on genetically defined cell identities and states.

## METHODS

### Genetic constructs

Mammalian codon-optimized MxEnc, MxB, MxC, MxD (UniProt: MXAN_3556, MXAN_3557, MXAN_4464, MXAN_2410), QtEnc (UniProt: A0A0F5HPP7_9BACI), QtIMEF (UniProt: A0A0F5HNH9_9BACI), MmZip14^FLAG^ (UniProt: Q75N73) and human H-chain ferritin (HHF) (UniProt: P02794) were custom synthesized by Integrated DNA Technologies and cloned into pcDNA3.1 (+) Zeocin (Invitrogen) using restriction cloning or Gibson assembly. A FLAG surface tag was C-terminally appended to MxEnc and QtEnc using Q5^®^ Site-Directed Mutagenesis (New England Biolabs). QtEnc^Farn^ was generated by appending a minimal C-terminal farnesylation signal (GCMSCKCVLS, 281-290 of rat H-Ras). Multigene expression of MxB, C and D was achieved by generating a single reading frame containing all three genes separated by P2A peptides yielding BCD_P2A_.

For a complete list of the genetic constructs featuring their composition refer to **Supplementary Table 1**.

### Cell culture, protein expression, and purification

Low passage number HEK293T (ECACC: 12022001) cells were cultured in advanced DMEM with 10 % FBS and Penicillin-Streptomycin at 100 μg/ml at 37 °C and 5 % CO_2_. Cells were transfected with X-tremeGENE HP (Roche) according to the protocol of the manufacturer. DNA amounts (ratio shell to cargo) were kept constant in all transient experiments to yield reproducible DNA-Lipoplex formation. To express combinations of MxEnc^FLAG^ or QtEnc^FLAG^ with cargo proteins, 70 % of the total DNA amount was encoding the shell and the remaining 30 % were used for the respective cargo molecule. For experiments with low-level Zip14 co-expression, the amount of shell was reduced to 65 % and 5 % of the total DNA amount was encoding Zip14. To facilitate iron loading, cells were supplemented with medium containing ferrous ammonium sulfate (FAS) at indicated concentrations 24 h after transfections. For ICP-MS analysis, encapsulins were purified from cells supplemented with 2 mM FAS.

For analysis of protein expression, cells were harvested between 24 and 48 h post-transfection and lysed with M-PER Mammalian Protein Extraction Reagent (Pierce Biotechnology) containing a mammalian protease inhibitor cocktail (SIGMA P8340, Sigma-Aldrich) according to the protocol of the manufacturer. After spinning down cell debris at 10,000 x g for 15 min, cell lysates were kept at 4 °C for downstream analyses. Protein concentrations of lysates were determined by measuring OD at 280 nm.

FLAG-tagged encapsulins were purified from HEK293T cells using the ANTI-FLAG M2 Affinity Gel (SIGMA A2220, Sigma-Aldrich) in a batch format according to the manufacturer’s protocol. In brief, cells were washed twice with DPBS and incubated with cold lysis buffer (50 mM Tris HCl, 150 mM NaCl, pH 7.4 + 1 % Triton X-100) for 15 min on a shaker. Cell debris was spun down at 10,000 x g for 15 min, and the supernatant was incubated with pre-equilibrated resin for 2 h at 4 °C. After protein binding, the flow-through was collected, and the resin washed with 5 column volumes (CV) TBS. For elution, the resin was incubated with 1 CV of 100 μg/ml 3x FLAG tag peptide (SIGMA F4799, Sigma-Aldrich) for 30 min at 4 °C. After the eluate was collected, the elution process was repeated once. Both eluate fractions were pooled and kept at 4 °C for further analysis. To analyze purified proteins, samples were mixed with SDS-PAGE sample buffer and incubated at 95 °C for 5 min. Samples were loaded onto pre-cast 12 % Bio-Rad Mini-PROTEAN^®^ TGX™ (Bio-Rad Laboratories) gels and run for 45 min at 200 V. Gels were either silver-stained using SilverQuest™ Silver Staining Kit (Novex) according to the protocol of the manufacturer or Coomassie stained using Bio-Safe™ Coomassie Stain (Bio-Rad Laboratories). For densitometric determination of SDS-PAGE gel bands, band intensity integrals of Coomassie-stained gels were measured using ImageJ (NIH).

### Blue Native gel electrophoresis and on-gel analyses

For detection of native encapsulins, pre-cast NativePAGE™ Novex^®^ 3–12 % Bis-Tris gels (Life Technologies) were used according to the manufacturer’s protocol. Gels were loaded with either FLAG-purified proteins or whole cell lysates mixed with NativePAGE™ Novex^®^ sample buffer and run for 120 min at 150 V. Unstained Protein Standard (Life Technologies) covering a size range between 20 - 1200 kDa was used as a marker. The total protein amount of whole cell lysates loaded per well was adjusted to ~1-3 μg.

Gels loaded with whole-cell lysates or purified samples were Coomassie-stained using Bio-Safe™ Coomassie Stain (Bio-Rad Laboratories). For detection of iron-containing proteins, gels were Prussian Blue (PB) stained. Briefly, gels were incubated in 2 % potassium hexacyanoferrate(II) in 10 % HCl for 45 min. For 3,3’-Diaminobenzidine (DAB)-enhancement (DAB PB), gels were washed three times with ddH_2_O and incubated in 0.1 M phosphate buffer pH 7.4 containing 0.025 % DAB and 0.005 % H_2_O_2_ until dark-brown bands appeared. To stop DAB polymerization, gels were washed three times with ddH_2_O.

### Dynamic Light Scattering

Data were obtained on a DynaPro NanoStar instrument and analyzed with DYNAMICS 7.1.9 software (Wyatt Technology). Measurements were performed at 22 °C using standard rectangular cuvettes containing 60 μl of protein sample in the concentration range between 0.15 and 0.5 mg/ml. For each measurement, 100 acquisitions with an acquisition time of 5 s were recorded.

### Quantification of cellular and protein iron content

To quantify the iron content of purified encapsulin proteins, inductively coupled plasma-mass spectrometry (ICP-MS) was performed. Measurements were performed on a NexION 350D (Perkin Elmer) in collision mode with kinetic energy discrimination (KED), and samples were diluted 1:1 with 65 % nitric acid and incubated at 65 °C overnight to release all iron.

To quantify the iron content of cell lysates, a commercial iron assay kit (Sigma) was used according to the user’s manual. In short, cleared supernatant was incubated with Iron Reducer for 30 min at RT to reduce Fe^3+^ to Fe^2+^ before adding Iron Probe. The reaction was incubated for 60 min at RT and absorbance was measured at 593 nm using a plate reader (SpectraMax M5, Molecular Devices). To calculate iron concentrations, a standard curve was plotted in the range of 0-14 μM iron.

### Cell viability assay

Iron-related cytotoxicity was monitored with a Luciferase-based viability assay (RealTime-Glo™ MT Cell Viability Assay, Promega) according to the protocol of the manufacturer in 96-well plate format as an endpoint measurement. Luminescence readings were taken on a Centro LB 960 (Berthold Technologies) at 0.5 s acquisition time.

### Cryo-electron Microscopy

Vitrified specimens of QtEnc were prepared in a Vitrobot (Thermofisher, Germany) at 22.0 °C and 90 % humidity. 3 μl of the particle dispersion was applied on 400-mesh C-Flat CF-2/1-4C copper grids (Protochips, USA) negatively glow discharged with a Plasma Cleaner (EMS, USA) at 50 mA for 45 s. After 15 s equilibration, the excess of liquid was removed through blotting (3 s, “force” −1) and the grids were automatically vitrified in liquid ethane below −172 °C. The samples were imaged using a Titan Krios microscope (Thermofisher, Germany), equipped with a Falcon 3EC direct electron detector. The system was operated at an acceleration voltage of 300 kV under low dose conditions (EPU acquisition software). Micrographs were collected in linear mode, for each of the three datasets. The dataset of QtEnc without cargo was acquired at a magnification of 37 kx (corresponding to a magnified pixel size of 1.8 Å), the total dose of 60 e-/Å^2^ was split along 13 frames and defocus values from −1 μm to −2 μm were applied. The two datasets from independently produced samples of QtEnc+QtIMEF were acquired at a magnification of 47 kx/37 kx (corresponding to a pixel size of 1.3/1.8Å) and the total dose of 60/40 e-/Å^2^ was split along 13/9 frames.

### Image processing and 3D reconstruction

All three datasets were processed using the RELION3.0^32^ image processing pipeline. 2D micrograph movies were motion corrected using the RELION3 MotionCor2 implementation. Contrast transfer function values were estimated with Ctffind4.1^33^. Particles were selected with Cryolo^34^ for QtEnc or manually for QtEnc+QtIMEF and subsequent reference-free 2D classification was performed with 2x downscaled particles in RELI0N3.0. This yielded 6836 particles for QtEnc and 362/518 particles for QtEnc+QtIMEF. For all datasets, an initial model with imposed I1 symmetry was generated inside RELION3.0 and subsequent 3D classification and 3D Auto-refinement resulted in the final maps. Postprocessing resulted in final resolutions of 7.6 Å for QtEnc and 13/11 Å for QtEnc+QtIMEF.

### Transmission Electron Microscopy

HEK cells were grown on aclar sheets (Science Services), fixed in 2.5 % glutaraldehyde (EM-grade, Science Services) and 0.05 % malachite green (Sigma Aldrich) in 0.1 M sodium cacodylate buffer (pH 7.4). After postfixation in 1 % osmium tetroxide, 0.8 % potassium ferrocyanide in 0.1 M sodium cacodylate buffer and a mordant step in 1 % tannic acid (Sigma Aldrich), cells were contrasted in 0.5 % uranyl acetate (Science Services). After dehydration, infiltration in epon (Serva) and curing, cell monolayers were ultrathin sectioned at 80 nm on formvar-coated copper grids (Plano) and postcontrasted using 1 % uranyl acetate and ultrostain (Leica). TEM images for **Fig. 4** and **Supplementary Fig. 5** were acquired on a JEM 1400plus (JEOL) using the TEMCenter software (JEOL).

For the electron microscopy data shown in **Supplementary Fig. 6**, cells were fixed in 2.5 % electron microscopy grade glutaraldehyde in 0.1 M sodium cacodylate buffer pH 7.4 (Science Services, Munich, Germany) and postfixed in 2 % aqueous osmium tetraoxide (Dalton, 1952). The samples were then dehydrated in gradual ethanol (30 - 100 %) and propylene oxide, embedded in Epon (Merck, Darmstadt, Germany) and cured for 24 h at 60 °C. Semithin sections were cut and stained with toluidine blue and sections of 50 nm were collected onto 200 mesh copper grids, stained with uranyl acetate and lead citrate before examination by transmission electron microscopy (Zeiss Libra 120 Plus, Carl Zeiss NTS GmbH, e3 Cell Metabolism 25, 1334–1347.e1–e4, June 6, 2017, Oberkochen, Germany). Pictures were acquired using a Slow Scan CCD-camera and iTEM software (Olympus Soft Imaging Solutions, Munster, Germany).

### Deep Learning assisted image classification

A dataset of 56 EM images of size 5000×5000 px was acquired to assess human and automatic classification performance. It was assured that each image only showed one type of nanospheres belonging to one of the three classes QtEnc+QtIMEF, MxEnc+MxBCD or ferritin. Due to the large image size and relatively small extent of ROIs containing the objects of interests, we adapted a pre-trained residual network^29^ as the backbone for dense feature extraction across the two image dimensions, yielding a 2048×157×157 feature tensor for each full-size image. On top, we implemented a gating mechanism to enable the possibility to focus the regions of interest and gain insights into the model’s decision process. The model was trained using a 5-fold cross-validation scheme with 45 training and 11 testing images per fold. For each location in the feature map, an object probability along with a class-probability was computed. The final class scores were determined by weighting class probabilities with object probabilities and summing over all pixels to arrive at a single score for the whole image, following

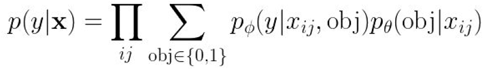

Implicitly, this model learns to simultaneously localize and classify regions of interest, using only global label information. No exhaustive hyperparameter search was performed. Deep learning models were trained using PyTorch^35^ and the ResNet50 implementation that is part of the torchvision library. For a detailed insight into the model architecture and parameters, we will release the source code under a permissive open source license. To estimate an upper bound of the possible performance, the same dataset used to train the model was blinded and distributed to three human labelers with domain knowledge (two of them were co-authors), who classified the images.

### Semantic Segmentation

Extending the previous experiments, objects in a subset of the dataset were exhaustively labeled using circular masks, yielding 400 individual annotations. An FCN with U-net shape^36^ was trained and evaluated on this dataset using 5-fold cross-validation over the full-sized images. The size of encapsulins expressed in cells was measured manually for 100 nanospheres using ImageJ. To generate a histogram of mean grey values, 266 encapsulin nanocompartments from 6 (QtEnc+QtIMEF) or 11 (MxEnc+MxBCD) TEM pictures were manually marked with ROIs and the histogram of each nanosphere was generated using ImageJ.

### Ferroxidase activity measurement

FLAG-tagged protein was purified from HEK293T cells expressing either QtEnc+QtIMEF or McEnc+MxBCD and protein concentrations were determined densitometrically via SDS-PAGE gel analysis. For the ferroxidase assay, 16 μM of QtIMEF or MxBCD inside the respective encapsulin shells were incubated with 50 Fe^2+^ per cargo in TBS buffer and the absorption of Fe^3+^ at 350 nm was monitored for 20 min in a plate reader (SpectraMax M5, Molecular Devices). For background correction, the absorption of free iron in buffer was recorded in parallel and subtracted from all measured values.

### Statistical analysis

One-way ANOVA, two-way ANOVA or paired t-tests followed by Bonferroni multiple comparison correction were performed using GraphPad Prism 6.05 (GraphPad Software, San Diego, California USA). All error bars given are average values ± SEM. A detailed table of all statistical analyses is given in **Supplementary Table 3**.

## ACKNOWLEDGMENTS

We are grateful for support from the European Research Council under grant agreements ERC-StG: 311552 (F.S., G.G.W.) and for support by Deutsche Forschungsgemeinschaft (DFG) through the TUM International Graduate School of Science and Engineering (IGSSE, project BIOMAG; G.G.W., S.P.). We thank Hannes Rolbieski for assistance with cell culture and the Simmel Laboratory for help with the DLS measurements. We thank the Institute of Hydrochemistry (TUM, Chair of Analytical Chemistry and Water Chemistry) for ICP-MS measurements. We thank Prof. Dr. Paola Turano and Dr. Silvia Ciambellotti for scientific advice on the ferroxidase measurements.

## AUTHOR CONTRIBUTIONS

F.S. and S.P. jointly conducted all molecular biology work and biochemical analyses and co-designed the experiments, F.Sch., M.K., and H.D. contributed the cryo-EM data and analysis. M.S. and T.M. generated cellular TEM data shown in Fig. 4. and Supplementary Fig. 5. M.A. and A.W. generated the TEM data shown in Supplementary Fig. 6. S.S. built the neural-net-based classification system. G.G.W. designed and supervised the study, F.S., S.P., and G.G.W. wrote the manuscript.

## DATA AVAILABILITY

Data are available from the corresponding author upon reasonable request.

## REFERENCES

1. Beck, M. & Baumeister, W. Cryo-Electron Tomography: Can it Reveal the Molecular Sociology of Cells in Atomic Detail? Trends Cell Biol. 26, 825–837 (2016).

2. Phillips, M. J. & Voeltz, G. K. Structure and function of ER membrane contact sites with other organelles. Nat. Rev. Mol. Cell Biol. 17, 69–82 (2016).

3. Denk, W., Briggman, K. L. & Helmstaedter, M. Structural neurobiology: missing link to a mechanistic understanding of neural computation. Nat. Rev. Neurosci. 13, 351–358 (2012).

4. Swanson, L. W. & Lichtman, J. W. From Cajal to Connectome and Beyond. Annu. Rev. Neurosci. 39, 197–216 (2016).

5. Martell, J. D. et al. Engineered ascorbate peroxidase as a genetically encoded reporter for electron microscopy. Nat. Biotechnol. 30, 1143–1148 (2012).

6. Lam, S. S. et al. Directed evolution of APEX2 for electron microscopy and proximity labeling. Nat. Methods 12, 51–54 (2015).

7. Shu, X. et al. A genetically encoded tag for correlated light and electron microscopy of intact cells, tissues, and organisms. PLoS Biol. 9, e1001041 (2011).

8. Joesch, M. et al. Reconstruction of genetically identified neurons imaged by serial-section electron microscopy. Elife 5, (2016).

9. Bouchet-Marquis, C., Pagratis, M., Kirmse, R. & Hoenger, A. Metallothionein as a clonable high-density marker for cryo-electron microscopy. J. Struct. Biol. 177, 119–127 (2012).

10. Hagiwara, H., Aoki, T., Suzuki, T. & Takata, K. Double-Label Immunoelectron Microscopy for Studying the Colocalization of Proteins in Cultured Cells. in Immunoelectron Microscopy: Methods and Protocols (eds. Schwartzbach, S. D. & Osafune, T.) 249–257 (Humana Press, 2010).

11. Fang, T. et al. Nanobody immunostaining for correlated light and electron microscopy with preservation of ultrastructure. Nat. Methods 15, 1029–1032 (2018).

12. Pallotto, M., Watkins, P. V., Fubara, B., Singer, J. H. & Briggman, K. L. Extracellular space preservation aids the connectomic analysis of neural circuits. Elife 4, (2015).

13. Shahidi, R. et al. A serial multiplex immunogold labeling method for identifying peptidergic neurons in connectomes. Elife 4, (2015).

14. Wang, Q., Mercogliano, C. P. & Löwe, J. A ferritin-based label for cellular electron cryotomography. Structure 19, 147–154 (2011).

15. Matsumoto, Y., Chen, R., Anikeeva, P. & Jasanoff, A. Engineering intracellular biomineralization and biosensing by a magnetic protein. Nat. Commun. 6, 8721 (2015).

16. Clarke, N. I. & Royle, S. J. FerriTag is a new genetically-encoded inducible tag for correlative light-electron microscopy. Nat. Commun. 9, 2604 (2018).

17. McHugh, C. A. et al. A virus capsid-like nanocompartment that stores iron and protects bacteria from oxidative stress. EMBO J. 33, 1896–1911 (2014).

18. Valdés-Stauber, N. & Scherer, S. Isolation and characterization of Linocin M18, a bacteriocin produced by Brevibacterium linens. Appl. Environ. Microbiol. 60, 3809–3814 (1994).

19. Sutter, M. et al. Structural basis of enzyme encapsulation into a bacterial nanocompartment. Nat. Struct. Mol. Biol. 15, 939–947 (2008).

20. Giessen, T. W. & Silver, P. A. Widespread distribution of encapsulin nanocompartments reveals functional diversity. Nat Microbiol 2, 17029 (2017).

21. Nichols, R. J., Cassidy-Amstutz, C., Chaijarasphong, T. & Savage, D. F. Encapsulins: molecular biology of the shell. Crit. Rev. Biochem. Mol. Biol. 52, 583–594 (2017).

22. Rurup, W. F., Snijder, J., Koay, M. S. T., Heck, A. J. R. & Cornelissen, J. J. L. M. Self-sorting of foreign proteins in a bacterial nanocompartment. J. Am. Chem. Soc. 136, 3828–3832 (2014).

23. Tamura, A. et al. Packaging guest proteins into the encapsulin nanocompartment from Rhodococcus erythropolis N771. Biotechnol. Bioeng. 112, 13–20 (2015).

24. Cassidy-Amstutz, C. et al. Identification of a Minimal Peptide Tag for in Vivo and in Vitro Loading of Encapsulin. Biochemistry 55, 3461–3468 (2016).

25. Putri, R. M. et al. Structural Characterization of Native and Modified Encapsulins as Nanoplatforms for in Vitro Catalysis and Cellular Uptake. ACS Nano 11, 12796–12804 (2017).

26. Sigmund, F. et al. Bacterial encapsulins as orthogonal compartments for mammalian cell engineering. Nat. Commun. 9, 1990 (2018).

27. Jenkitkasemwong, S., Wang, C.-Y., Mackenzie, B. & Knutson, M. D. Physiologic implications of metal-ion transport by ZIP14 and ZIP8. Biometals 25, 643–655 (2012).

28. Bernacchioni, C., Pozzi, C., Di Pisa, F., Mangani, S. & Turano, P. Ferroxidase Activity in Eukaryotic Ferritin is Controlled by Accessory-Iron-Binding Sites in the Catalytic Cavity. Chemistry 22, 16213–16219 (2016).

29. He, K., Zhang, X., Ren, S. & Sun, J. Deep Residual Learning for Image Recognition. arXiv [cs.CV] (2015).

30. Apolloni, A., Prior, I. A., Lindsay, M., Parton, R. G. & Hancock, J. F. H-ras but not K-ras traffics to the plasma membrane through the exocytic pathway. Mol. Cell. Biol. 20, 2475–2487 (2000).

31. He, D. et al. Structural characterization of encapsulated ferritin provides insight into iron storage in bacterial nanocompartments. Elife 5, (2016).

32. Zivanov, J. et al. New tools for automated high-resolution cryo-EM structure determination in RELION-3. Elife 7, (2018).

33. Rohou, A. & Grigorieff, N. CTFFIND4: Fast and accurate defocus estimation from electron micrographs. J. Struct. Biol. 192, 216–221 (2015).

34. Wagner, T. et al. SPHIRE-crYOLO: A fast and well-centering automated particle picker for cryo-EM. bioRxiv 356584 (2018). doi:10.1101/356584

35. Paszke, A., Gross, S. & Chintala, S. Automatic differentiation in PyTorch.

36. Ronneberger, O., Fischer, P. & Brox, T. U-Net: Convolutional Networks for Biomedical Image Segmentation. arXiv [cs.CV] (2015).

