## Supplementary Information for "Iron-sequestering nanocompartments as multiplexed Electron Microscopy gene reporters"

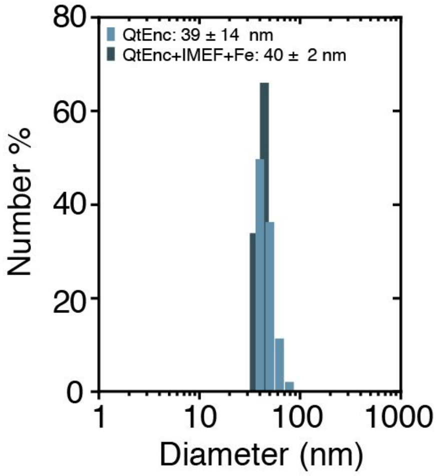


**Supplementary Fig. 1 | DLS measurement of purified encapsulin nanospheres.** Dynamic light scattering (DLS) measurement of purified QtEnc without or with QtIMEF and iron loading purified from HEK293T cells. The histogram shows monodisperse distributions for both samples.

* contributed equally
† corresponding author

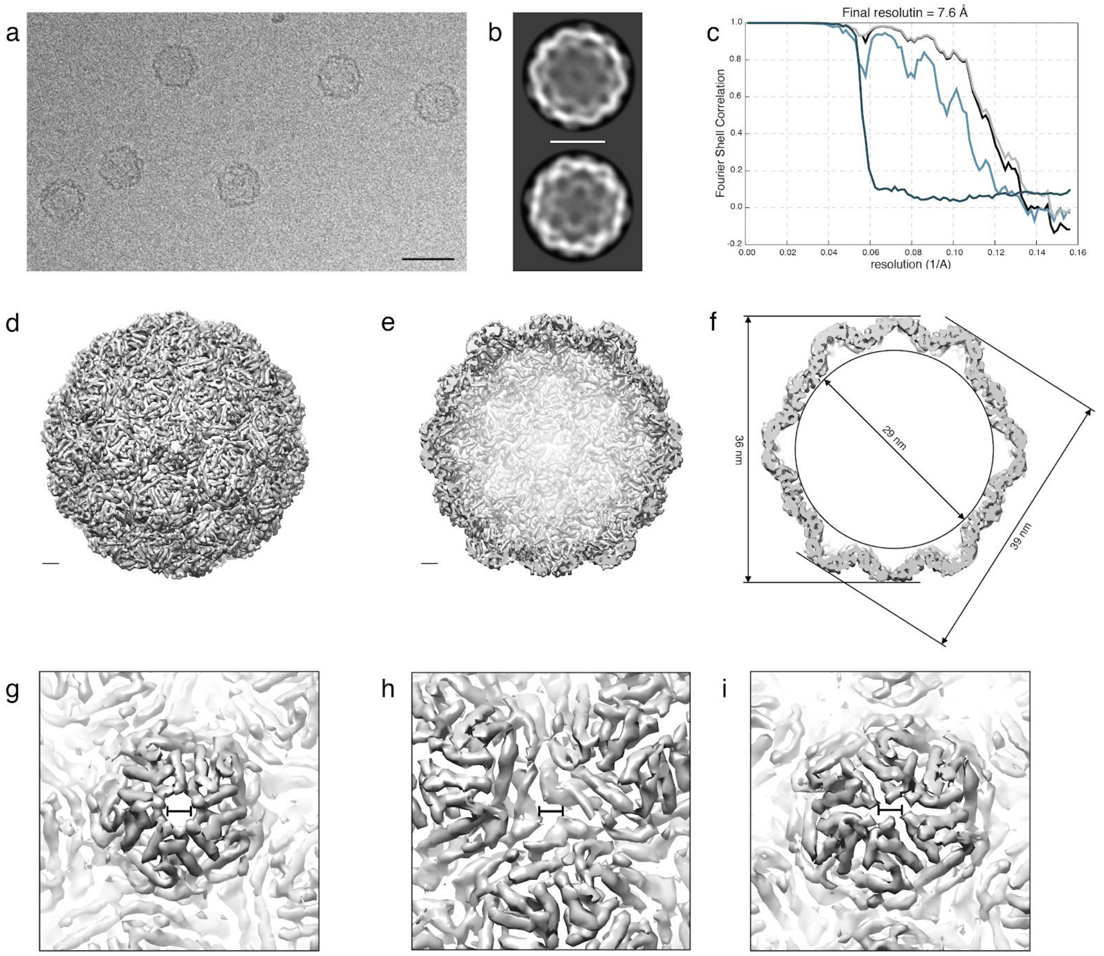


**Supplementary Fig. 2 | Cryo-EM of QtEnc nanocompartments.** (**a**) Exemplary micrograph of QtEnc nanospheres purified from mammalian cells without co-expressed cargo. Scale bar represents 50 nm. (**b)** Exemplary 2D class averages of QtEnc; scale bar is 25 nm. (**c**) FSC curves of the final post-processed map showing the Fourier shell correlations of unmasked (light turquoise) and masked (light grey) maps as well as corrected curve (black) and corrected curve of phase randomized masked maps (dark turquoise). (**d)** Electron density map of QtEnc; scale bar represents 2 nm. (**e**) Cutaway view of the QtEnc shell without cargo; scale bar is 2 nm. (**f)** Inner and outer diameters (through twofold and fivefold axes) shown on a slice representation through the center of QtEnc without cargo. (**g**) Close-up of the fivefold symmetry center revealing a putative pore region with a diameter of about ~1 nm. (**h)** Zoomed-in view of the threefold symmetry center with an electron-sparse region in its center. (**i**) Zoom-in to the twofold symmetry center showing a cleft-like electron sparse region. Scale bars in g,h,i represent 1 nm.

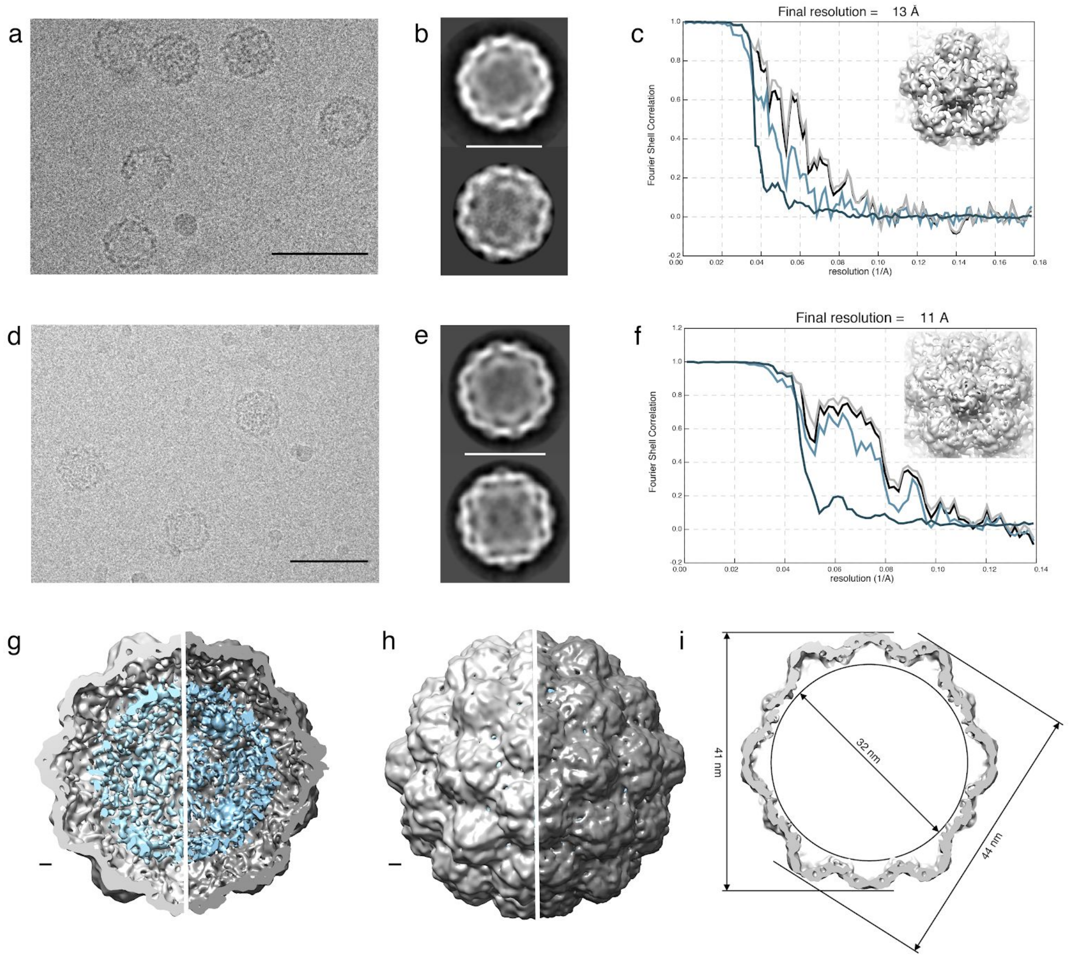


**Supplementary Fig. 3 | Cryo-EM of QtEnc with QtIMEF.** (**a,d**) Exemplary micrographs of two independent samples from QtEnc+QtIMEF purified from mammalian cells. Scale bars represent 48 nm. (**b,e**) Exemplary 2D class averages of QtEnc with QtIMEF; scale bars represent 24 nm. (**c,f**) FSC curves of the final post-processed map showing the Fourier shell correlations of unmasked (light turquoise) and masked (light grey) maps as well as corrected curve (black) and corrected curve of phase randomized masked maps (dark turquoise). (**g,h**) Electron density maps of QtEnc+QtIMEF as cutaway and surface view. Light grey halves are reconstructed from the sample shown in a,b,c and dark gray halves from samples shown in d,e,f. The cargo and shell are shown at different electron density thresholds for each sample. Scale bar represents 2 nm. (**i**) Inner and outer diameters (through twofold and fivefold axes) shown on a slice representation through the center of the loaded QtEnc (cargo not shown).

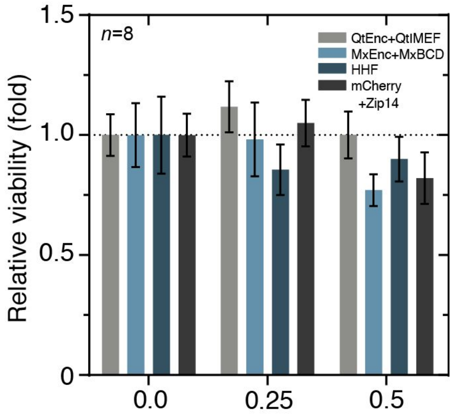


**Supplementary Fig. 4 | Luciferase-based viability assay of HEK293T cells expressing QtEnc+QtIMEF, MxEnc+MxBCD, HHF or mCherry.** In all conditions, low-level Zip14 was co-expressed to boost ferrous iron uptake, which was supplemented at different concentrations (0-0.5 mM) for 36 h prior to analysis. Bars show the mean ± SEM of *n*=8 independent biological replicates. Luciferase signals were normalized to those obtained for cells without FAS supplementation to obtain a measure of relative viability. 2-way ANOVA resulted in non-significant main effects for FAS concentration ($p$=0.1834), gene type ($p$=0.4947) and interaction ($p$=0.7883). For a complete list of post hoc test results please see Supplementary Table 3.

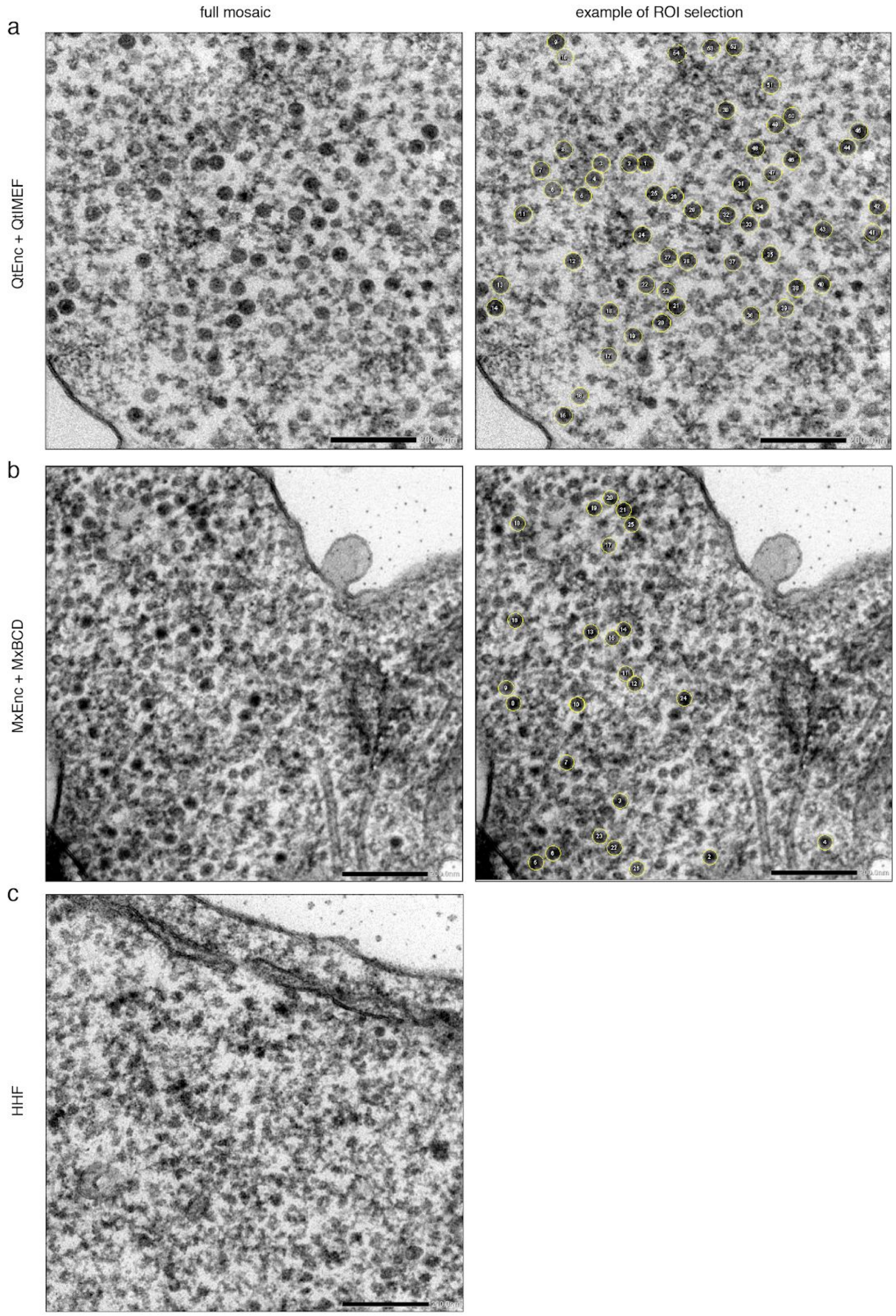


**Supplementary Fig. 5 | Manual segmentation of intracellular encapsulins in TEM.** Exemplary full frame EM image of HEK293T cells expressing QtEnc+QtIMEF (**a**) or MxEnc+MxBCD (**b**) together with low-level Zip14 that were supplemented with 0.5 mM FAS for 36 h before fixation. Exemplary images on the right indicate manually segmented ROIs (yellow circles) used to construct the histograms shown in Fig 4c. (**c**) Exemplary full frame EM image of HEK293T cells expressing human H-chain ferritin (HHF). ROI selection was not performed for this condition because ferritin particles could not be differentiated from ribosomes and other endogenous cellular structures. Scale bars represent 200 nm.

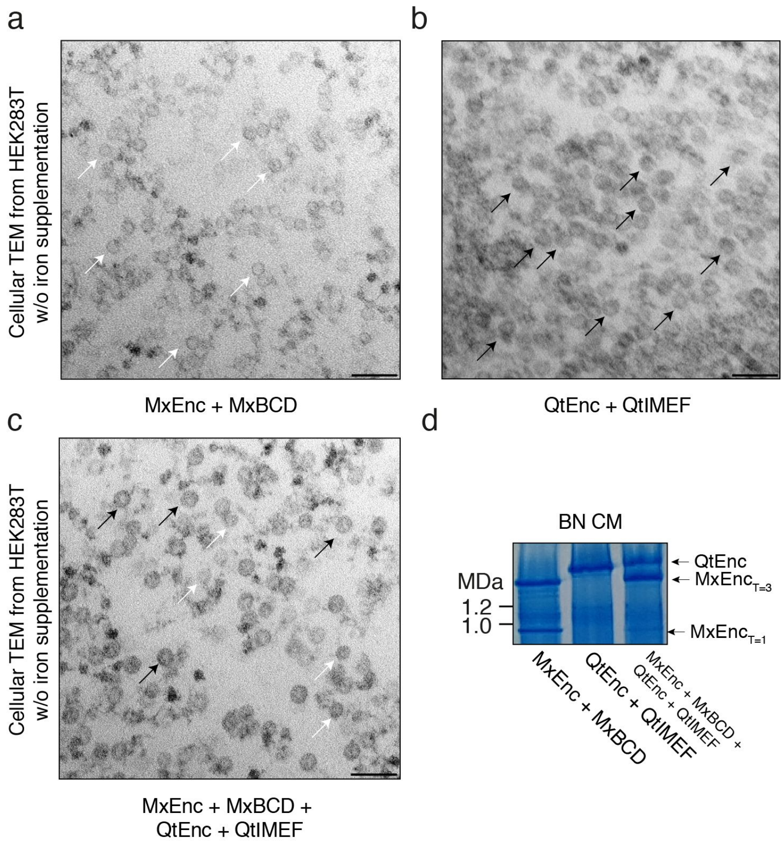


**Supplementary Fig. 6 | TEM of HEK293T cells expressing encapsulins without iron supplementation. (a)** TEM image of HEK293T cells expressing MxEnc+MxBCD without iron supplementation. White arrows indicate Mx encapsulins. (**b)** Cellular TEM image of HEK293T cells expressing QtEnc+QtIMEF. Black arrows indicate Qt encapsulins. (**c)** TEM image of HEK293T cells expressing a mixture of MxEnc+MxBCD and QtEnc+QtIMEF. Assembled encapsulins of both sizes are apparent. White arrows indicate Mx encapsulins (31.8 ± 2.6 nm) and black arrows indicate Qt encapsulins (38.7 ± 2.3 nm). TEM images were acquired on a Zeiss Libra 120 Plus. Scale bar in all images represents 100 nm. (**d)** Exemplary Coomassie-stained BN-PAGE (BN CM) loaded with whole cell lysates of HEK293T expressing the gene combinations as shown in the EM images. The two distinct bands in the right lane indicate no intermixing of monomeric subunits upon co-expression of both encapsulin:cargo systems.

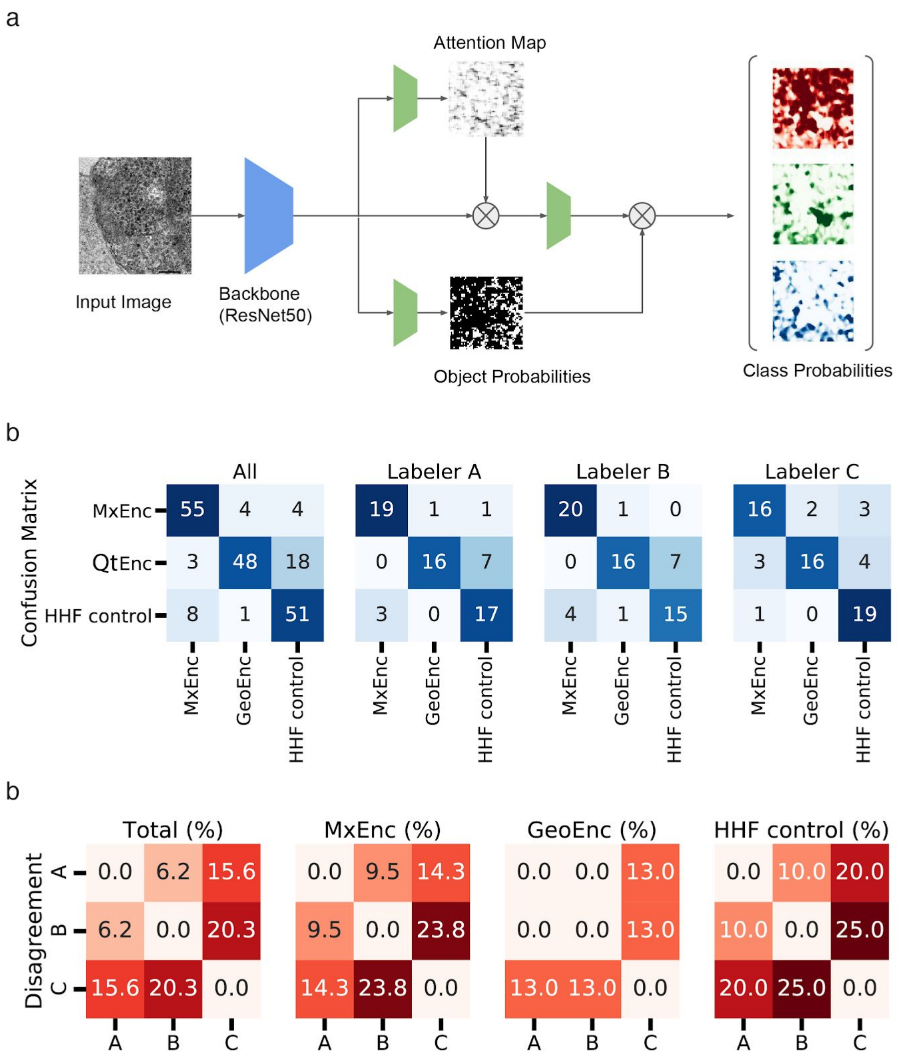


**Supplementary Fig. 7 | Deep learning approach for classification of TEM images. (a)** Deep learning approach using a ResNet50 with added point-wise attention modules for classification of TEM images. Attention modules allow the network to base the final prediction only on specific parts of the image that are relevant for the differentiation of the three classes. (**b)** Confusion matrices showing the qualitative results of TEM image classification for different human labelers. (**c**) Confusion matrix showing the quantitative analysis of image classification for each class of iron-storing proteins as performed by humans (*n*=3 individual classifications).

**Supplementary Table 1 | Complete list of genetic constructs.** Expression constructs encoding variants of the encapsulin shells and different cargo proteins are specified.

| **Encapsulin shell proteins** | | |
|---|---|---|
| pcDNA 3.1 (+) Zeocin MxEncA$^{FLAG}$ | MxEncA-GSG-DYKDDDDK* | UniProt: EncA: MXAN_3556 |
| pcDNA 3.1 (+) Zeocin QtEnc$^{FLAG}$ | QtEnc-GSG-DYKDDDDK* | WP_039238471.1, UniProt: A0A0F5HPP7_9BACI |
| pcDNA 3.1 (+) Zeocin QtEnc$^{Farn}$ | QtEnc-GSG-GCMSCKCVLS* | WP_039238471.1, UniProt: A0A0F5HPP7_9BACI |
| **Encapsulin cargo proteins** | | |
| pcDNA 3.1 (+) Zeocin MxEncBCD$^{P2A}$ | MxEncB-GSGATNFSLLKQAGDVEENPGP-MxEncC-GSGATNFSLLKQAGDVEENPGP-MxEncD* | UniProt: EncB: MXAN_3557, EncC: MXAN_4464, EncD: MXAN_2410 |
| pcDNA 3.1 (+) Zeocin QtIMEF | QtIMEF* | WP_039238473.1, UniProt: A0A0F5HNH9_9BACI |
| **Other proteins** | | |
| pcDNA 3.1 (+) MmZip14-FLAG | MmZip14-GGGGGSGGGGS-DYKDDDDK* | UniProt: Q75N73 |
| pcDNA 3.1 (+) HHF | HHF* | UniProt: P02794 |

**Supplementary Table 2 | Properties of the two encapsulin systems.**

| | **MxEnc** | **QtEnc** |
|---|---|---|
| Symmetry of icosahedral shell | T=1 and T=3 | T=4 |
| Number of subunits | 60 (T=1), 180 (T=3) | 240 (T=4) |
| Molecular weight of monomers (kDa) [1] | MxEnc: 33; MxB: 17; MxC: 13; MxD: 11 | QtEnc: 32; QtIMEF: 23 |
| Number of cargo proteins per shell [2] | 86 ± 3 (MxB), 93 ± 9 (MxC), 50 ± 15 (MxD) | 231 ± 5 (QtIMEF) |
| Shell outer diameter of cargo loaded compartment (nm) [3] | ~32 | ~44 |
| Iron atoms per protein shell [4] | 18856 ± 807 | 35097 ± 853 |

[1] MxEnc: taken from reference [1], QtEnc: taken from reference[2]
[2] calculated from densitometric SDS-PAGE analysis from *n*=3 independent samples
[3] values for MxEnc (T=3 assembly) were determined using single particle cryoEM[1,3].
[4] measured by ICP-MS (*n*=2)

**Supplementary Table 3 | Results of the post hoc tests.**

| **Iron loading in mammalian cells without Zip14** | | | | |
|---|---|---|---|---|
| 0:McEnc+MxBCD vs. 0:QtEnc+IMEF | Two-way ANOVA, Bonferroni-correction | > 0.9999 | ns | n =3 |
| 0:McEnc+MxBCD vs. 0.125:McEnc+MxBCD | Two-way ANOVA, Bonferroni-correction | > 0.9999 | ns | n =3 |
| 0:McEnc+MxBCD vs. 0.125:QtEnc+IMEF | Two-way ANOVA, Bonferroni-correction | > 0.9999 | ns | n =3 |
| 0:McEnc+MxBCD vs. 0.25:McEnc+MxBCD | Two-way ANOVA, Bonferroni-correction | > 0.9999 | ns | n =3 |
| 0:McEnc+MxBCD vs. 0.25:QtEnc+IMEF | Two-way ANOVA, Bonferroni-correction | > 0.9999 | ns | n =3 |
| 0:McEnc+MxBCD vs. 0.50:McEnc+MxBCD | Two-way ANOVA, Bonferroni-correction | 0.,5833 | ns | n =3 |
| 0:McEnc+MxBCD vs. 0.50:QtEnc+IMEF | Two-way ANOVA, Bonferroni-correction | 0.0001 | *** | n =3 |
| 0:McEnc+MxBCD vs. 1:McEnc+MxBCD | Two-way ANOVA, Bonferroni-correction | 0.0036 | ** | n =3 |
| 0:McEnc+MxBCD vs. 1:QtEnc+IMEF | Two-way ANOVA, Bonferroni-correction | < 0.0001 | **** | n =3 |
| 0:QtEnc+IMEF vs. 0.125:McEnc+MxBCD | Two-way ANOVA, Bonferroni-correction | > 0.9999 | ns | n =3 |
| 0:QtEnc+IMEF vs. 0.125:QtEnc+IMEF | Two-way ANOVA, Bonferroni-correction | > 0.9999 | ns | n =3 |
| 0:QtEnc+IMEF vs. 0.25:McEnc+MxBCD | Two-way ANOVA, Bonferroni-correction | > 0.9999 | ns | n =3 |
| 0:QtEnc+IMEF vs. 0.25:QtEnc+IMEF | Two-way ANOVA, Bonferroni-correction | > 0.9999 | ns | n =3 |
| 0:QtEnc+IMEF vs. 0.50:McEnc+MxBCD | Two-way ANOVA, Bonferroni-correction | > 0.9999 | ns | n =3 |
| 0:QtEnc+IMEF vs. 0.50:QtEnc+IMEF | Two-way ANOVA, Bonferroni-correction | 0.0166 | * | n =3 |
| 0:QtEnc+IMEF vs. 1:McEnc+MxBCD | Two-way ANOVA, Bonferroni-correction | 0.4837 | ns | n =3 |
| 0:QtEnc+IMEF vs. 1:QtEnc+IMEF | Two-way ANOVA, Bonferroni-correction | < 0.0001 | **** | n =3 |
| 0.125:McEnc+MxBCD vs. 0.125:QtEnc+IMEF | Two-way ANOVA, Bonferroni-correction | > 0.9999 | ns | n =3 |
| 0.125:McEnc+MxBCD vs. 0.25:McEnc+MxBCD | Two-way ANOVA, Bonferroni-correction | > 0.9999 | ns | n =3 |
| 0.125:McEnc+MxBCD vs. 0.25:QtEnc+IMEF | Two-way ANOVA, Bonferroni-correction | > 0.9999 | ns | n =3 |
| 0.125:McEnc+MxBCD vs. 0.50:McEnc+MxBCD | Two-way ANOVA, Bonferroni-correction | > 0.9999 | ns | n =3 |

| | | | | |
|---|---|---|---|---|
| 0.125:McEnc+MxBCD vs. 0.50:QtEnc+IMEF | Two-way ANOVA, Bonferroni-correction | 0.0427 | * | n =3 |
| 0.125:McEnc+MxBCD vs. 1:McEnc+MxBCD | Two-way ANOVA, Bonferroni-correction | 0.8393 | ns | n =3 |
| 0.125:McEnc+MxBCD vs. 1:QtEnc+IMEF | Two-way ANOVA, Bonferroni-correction | < 0.0001 | **** | n =3 |
| 0.125:QtEnc+IMEF vs. 0.25:McEnc+MxBCD | Two-way ANOVA, Bonferroni-correction | > 0.9999 | ns | n =3 |
| 0.125:QtEnc+IMEF vs. 0.25:QtEnc+IMEF | Two-way ANOVA, Bonferroni-correction | > 0.9999 | ns | n =3 |
| 0.125:QtEnc+IMEF vs. 0.50:McEnc+MxBCD | Two-way ANOVA, Bonferroni-correction | > 0.9999 | ns | n =3 |
| 0.125:QtEnc+IMEF vs. 0.50:QtEnc+IMEF | Two-way ANOVA, Bonferroni-correction | 0.0012 | ** | n =3 |
| 0.125:QtEnc+IMEF vs. 1:McEnc+MxBCD | Two-way ANOVA, Bonferroni-correction | 0.0347 | * | n =3 |
| 0.125:QtEnc+IMEF vs. 1:QtEnc+IMEF | Two-way ANOVA, Bonferroni-correction | < 0.0001 | **** | n =3 |
| 0.25:McEnc+MxBCD vs. 0.25:QtEnc+IMEF | Two-way ANOVA, Bonferroni-correction | > 0.9999 | ns | n =3 |
| 0.25:McEnc+MxBCD vs. 0.50:McEnc+MxBCD | Two-way ANOVA, Bonferroni-correction | > 0.9999 | ns | n =3 |
| 0.25:McEnc+MxBCD vs. 0.50:QtEnc+IMEF | Two-way ANOVA, Bonferroni-correction | 0.0073 | ** | n =3 |
| 0.25:McEnc+MxBCD vs. 1:McEnc+MxBCD | Two-way ANOVA, Bonferroni-correction | 0.2175 | ns | n =3 |
| 0.25:McEnc+MxBCD vs. 1:QtEnc+IMEF | Two-way ANOVA, Bonferroni-correction | < 0.0001 | **** | n =3 |
| 0.25:QtEnc+IMEF vs. 0.50:McEnc+MxBCD | Two-way ANOVA, Bonferroni-correction | > 0.9999 | ns | n =3 |
| 0.25:QtEnc+IMEF vs. 0.50:QtEnc+IMEF | Two-way ANOVA, Bonferroni-correction | 0.0249 | * | n =3 |
| 0.25:QtEnc+IMEF vs. 1:McEnc+MxBCD | Two-way ANOVA, Bonferroni-correction | 0.7104 | ns | n =3 |
| 0.25:QtEnc+IMEF vs. 1:QtEnc+IMEF | Two-way ANOVA, Bonferroni-correction | < 0.0001 | **** | n =3 |
| 0.50:McEnc+MxBCD vs. 0.50:QtEnc+IMEF | Two-way ANOVA, Bonferroni-correction | 0.0618 | ns | n =3 |
| 0.50:McEnc+MxBCD vs. 1:McEnc+MxBCD | Two-way ANOVA, Bonferroni-correction | > 0.,9999 | ns | n =3 |
| 0.50:McEnc+MxBCD vs. 1:QtEnc+IMEF | Two-way ANOVA, Bonferroni-correction | < 0.0001 | **** | n =3 |
| 0.50:QtEnc+IMEF vs. 1:McEnc+MxBCD | Two-way ANOVA, Bonferroni-correction | >0.9999 | ns | n =3 |

| | | | | |
|---|---|---|---|---|
| 0.50:QtEnc+IMEF vs. 1:QtEnc+IMEF | Two-way ANOVA, Bonferroni-correction | < 0.0001 | **** | n =3 |
| 1:McEnc+MxBCD vs. 1:QtEnc+IMEF | Two-way ANOVA, Bonferroni-correction | < 0.0001 | **** | n =3 |
| **Iron loading in mammalian cells with Zip14** | | | | |
| 0:McEnc+MxBCD vs. 0:QtEnc+IMEF | Two-way ANOVA, Bonferroni-correction | > 0.9999 | ns | n =3 |
| 0:McEnc+MxBCD vs. 0.125:McEnc+MxBCD | Two-way ANOVA, Bonferroni-correction | > 0.9999 | ns | n =3 |
| 0:McEnc+MxBCD vs. 0.125:QtEnc+IMEF | Two-way ANOVA, Bonferroni-correction | 0.1028 | ns | n =3 |
| 0:McEnc+MxBCD vs. 0.25:McEnc+MxBCD | Two-way ANOVA, Bonferroni-correction | 0.0105 | * | n =3 |
| 0:McEnc+MxBCD vs. 0.25:QtEnc+IMEF | Two-way ANOVA, Bonferroni-correction | <0.0001 | **** | n =3 |
| 0:McEnc+MxBCD vs. 0.50:McEnc+MxBCD | Two-way ANOVA, Bonferroni-correction | <0.0001 | **** | n =3 |
| 0:McEnc+MxBCD vs. 0.50:QtEnc+IMEF | Two-way ANOVA, Bonferroni-correction | < 0.0001 | **** | n =3 |
| 0:McEnc+MxBCD vs. 1:McEnc+MxBCD | Two-way ANOVA, Bonferroni-correction | < 0.0001 | **** | n =3 |
| 0:McEnc+MxBCD vs. 1:QtEnc+IMEF | Two-way ANOVA, Bonferroni-correction | < 0.0001 | **** | n =3 |
| 0:QtEnc+IMEF vs. 0.125:McEnc+MxBCD | Two-way ANOVA, Bonferroni-correction | > 0.9999 | ns | n =3 |
| 0:QtEnc+IMEF vs. 0.125:QtEnc+IMEF | Two-way ANOVA, Bonferroni-correction | 0.7190 | ns | n =3 |
| 0:QtEnc+IMEF vs. 0.25:McEnc+MxBCD | Two-way ANOVA, Bonferroni-correction | 0.0784 | ns | n =3 |
| 0:QtEnc+IMEF vs. 0.25:QtEnc+IMEF | Two-way ANOVA, Bonferroni-correction | < 0.0001 | **** | n =3 |
| 0:QtEnc+IMEF vs. 0.50:McEnc+MxBCD | Two-way ANOVA, Bonferroni-correction | 0.0001 | *** | n =3 |
| 0:QtEnc+IMEF vs. 0.50:QtEnc+IMEF | Two-way ANOVA, Bonferroni-correction | < 0.0001 | **** | n =3 |
| 0:QtEnc+IMEF vs. 1:McEnc+MxBCD | Two-way ANOVA, Bonferroni-correction | < 0.0001 | **** | n =3 |
| 0:QtEnc+IMEF vs. 1:QtEnc+IMEF | Two-way ANOVA, Bonferroni-correction | < 0.0001 | **** | n =3 |
| 0.125:McEnc+MxBCD vs. 0.125:QtEnc+IMEF | Two-way ANOVA, Bonferroni-correction | > 0.9999 | ns | n =3 |
| 0.125:McEnc+MxBCD vs. 0.25:McEnc+MxBCD | Two-way ANOVA, Bonferroni-correction | 0.1767 | ns | n =3 |

| | | | | |
|---|---|---|---|---|
| 0.125:McEnc+MxBCD vs. 0.25:QtEnc+IMEF | Two-way ANOVA, Bonferroni-correction | 0.0002 | *** | n =3 |
| 0.125:McEnc+MxBCD vs. 0.50:McEnc+MxBCD | Two-way ANOVA, Bonferroni-correction | 0.0003 | *** | n =3 |
| 0.125:McEnc+MxBCD vs. 0.50:QtEnc+IMEF | Two-way ANOVA, Bonferroni-correction | < 0.0001 | **** | n =3 |
| 0.125:McEnc+MxBCD vs. 1:McEnc+MxBCD | Two-way ANOVA, Bonferroni-correction | < 0.0001 | **** | n =3 |
| 0.125:McEnc+MxBCD vs. 1:QtEnc+IMEF | Two-way ANOVA, Bonferroni-correction | < 0.0001 | **** | n =3 |
| 0.125:QtEnc+IMEF vs. 0.25:McEnc+MxBCD | Two-way ANOVA, Bonferroni-correction | > 0.9999 | ns | n =3 |
| 0.125:QtEnc+IMEF vs. 0.25:QtEnc+IMEF | Two-way ANOVA, Bonferroni-correction | 0.0272 | * | n =3 |
| 0.125:QtEnc+IMEF vs. 0.50:McEnc+MxBCD | Two-way ANOVA, Bonferroni-correction | 0.0540 | ns | n =3 |
| 0.125:QtEnc+IMEF vs. 0.50:QtEnc+IMEF | Two-way ANOVA, Bonferroni-correction | < 0.0001 | **** | n =3 |
| 0.125:QtEnc+IMEF vs. 1:McEnc+MxBCD | Two-way ANOVA, Bonferroni-correction | < 0.0001 | **** | n =3 |
| 0.125:QtEnc+IMEF vs. 1:QtEnc+IMEF | Two-way ANOVA, Bonferroni-correction | < 0.0001 | **** | n =3 |
| 0.25:McEnc+MxBCD vs. 0.25:QtEnc+IMEF | Two-way ANOVA, Bonferroni-correction | 0.2626 | ns | n =3 |
| 0.25:McEnc+MxBCD vs. 0.50:McEnc+MxBCD | Two-way ANOVA, Bonferroni-correction | 0.5066 | ns | n =3 |
| 0.25:McEnc+MxBCD vs. 0.50:QtEnc+IMEF | Two-way ANOVA, Bonferroni-correction | < 0.0001 | **** | n =3 |
| 0.25:McEnc+MxBCD vs. 1:McEnc+MxBCD | Two-way ANOVA, Bonferroni-correction | < 0.0001 | **** | n =3 |
| 0.25:McEnc+MxBCD vs. 1:QtEnc+IMEF | Two-way ANOVA, Bonferroni-correction | < 0.0001 | **** | n =3 |
| 0.25:QtEnc+IMEF vs. 0.50:McEnc+MxBCD | Two-way ANOVA, Bonferroni-correction | >0.9999 | ns | n =3 |
| 0.25:QtEnc+IMEF vs. 0.50:QtEnc+IMEF | Two-way ANOVA, Bonferroni-correction | 0.0095 | ** | n =3 |
| 0.25:QtEnc+IMEF vs. 1:McEnc+MxBCD | Two-way ANOVA, Bonferroni-correction | <0.0001 | **** | n =3 |
| 0.25:QtEnc+IMEF vs. 1:QtEnc+IMEF | Two-way ANOVA, Bonferroni-correction | < 0.0001 | **** | n =3 |
| 0.50:McEnc+MxBCD vs. 0.50:QtEnc+IMEF | Two-way ANOVA, Bonferroni-correction | 0.0048 | ** | n =3 |
| 0.50:McEnc+MxBCD vs. 1:McEnc+MxBCD | Two-way ANOVA, Bonferroni-correction | < 0.0001 | **** | n =3 |

| | | | | |
|---|---|---|---|---|
| 0.50:McEnc+MxBCD vs. 1:QtEnc+IMEF | Two-way ANOVA, Bonferroni-correction | < 0.0001 | **** | n =3 |
| 0.50:QtEnc+IMEF vs. 1:McEnc+MxBCD | Two-way ANOVA, Bonferroni-correction | 0.0017 | ** | n =3 |
| 0.50:QtEnc+IMEF vs. 1:QtEnc+IMEF | Two-way ANOVA, Bonferroni-correction | 0.0001 | *** | n =3 |
| 1:McEnc+MxBCD vs. 1:QtEnc+IMEF | Two-way ANOVA, Bonferroni-correction | > 0.9999 | ns | n =3 |
| **Viability assay** | | | | |
| 0.00:mCherry vs. 0.00:HHF | Two-way ANOVA, Bonferroni-correction | p > 0.9999 | ns | n = 8 |
| 0.00:mCherry vs. 0.00:Mx | Two-way ANOVA, Bonferroni-correction | p > 0.9999 | ns | n = 8 |
| 0.00:mCherry vs. 0.00:Geo | Two-way ANOVA, Bonferroni-correction | p > 0.9999 | ns | n = 8 |
| 0.00:mCherry vs. 0.25:mCherry | Two-way ANOVA, Bonferroni-correction | p > 0.9999 | ns | n = 8 |
| 0.00:mCherry vs. 0.25:HHF | Two-way ANOVA, Bonferroni-correction | p > 0.9999 | ns | n = 8 |
| 0.00:mCherry vs. 0.25:Mx | Two-way ANOVA, Bonferroni-correction | p > 0.9999 | ns | n = 8 |
| 0.00:mCherry vs. 0.25:Geo | Two-way ANOVA, Bonferroni-correction | p > 0.9999 | ns | n = 8 |
| 0.00:mCherry vs. 0.50:mCherry | Two-way ANOVA, Bonferroni-correction | p > 0.9999 | ns | n = 8 |
| 0.00:mCherry vs. 0.50:HHF | Two-way ANOVA, Bonferroni-correction | p > 0.9999 | ns | n = 8 |
| 0.00:mCherry vs. 0.50:Mx | Two-way ANOVA, Bonferroni-correction | p > 0.9999 | ns | n = 8 |
| 0.00:mCherry vs. 0.50:Geo | Two-way ANOVA, Bonferroni-correction | p > 0.9999 | ns | n = 8 |
| 0.00:HHF vs. 0.00:Mx | Two-way ANOVA, Bonferroni-correction | p > 0.9999 | ns | n = 8 |
| 0.00:HHF vs. 0.00:Geo | Two-way ANOVA, Bonferroni-correction | p > 0.9999 | ns | n = 8 |
| 0.00:HHF vs. 0.25:mCherry | Two-way ANOVA, Bonferroni-correction | p > 0.9999 | ns | n = 8 |
| 0.00:HHF vs. 0.25:HHF | Two-way ANOVA, Bonferroni-correction | p > 0.9999 | ns | n = 8 |
| 0.00:HHF vs. 0.25:Mx | Two-way ANOVA, Bonferroni-correction | p > 0.9999 | ns | n = 8 |
| 0.00:HHF vs. 0.25:Geo | Two-way ANOVA, Bonferroni-correction | p > 0.9999 | ns | n = 8 |

| | | | | |
|---|---|---|---|---|
| 0.00:HHF vs. 0.50:mCherry | Two-way ANOVA, Bonferroni-correction | p > 0.9999 | ns | n = 8 |
| 0.00:HHF vs. 0.50:HHF | Two-way ANOVA, Bonferroni-correction | p > 0.9999 | ns | n = 8 |
| 0.00:HHF vs. 0.50:Mx | Two-way ANOVA, Bonferroni-correction | p > 0.9999 | ns | n = 8 |
| 0.00:HHF vs. 0.50:Geo | Two-way ANOVA, Bonferroni-correction | p > 0.9999 | ns | n = 8 |
| 0.00:Mx vs. 0.00:Geo | Two-way ANOVA, Bonferroni-correction | p > 0.9999 | ns | n = 8 |
| 0.00:Mx vs. 0.25:mCherry | Two-way ANOVA, Bonferroni-correction | p > 0.9999 | ns | n = 8 |
| 0.00:Mx vs. 0.25:HHF | Two-way ANOVA, Bonferroni-correction | p > 0.9999 | ns | n = 8 |
| 0.00:Mx vs. 0.25:Mx | Two-way ANOVA, Bonferroni-correction | p > 0.9999 | ns | n = 8 |
| 0.00:Mx vs. 0.25:Geo | Two-way ANOVA, Bonferroni-correction | p > 0.9999 | ns | n = 8 |
| 0.00:Mx vs. 0.50:mCherry | Two-way ANOVA, Bonferroni-correction | p > 0.9999 | ns | n = 8 |
| 0.00:Mx vs. 0.50:HHF | Two-way ANOVA, Bonferroni-correction | p > 0.9999 | ns | n = 8 |
| 0.00:Mx vs. 0.50:Mx | Two-way ANOVA, Bonferroni-correction | p > 0.9999 | ns | n = 8 |
| 0.00:Mx vs. 0.50:Geo | Two-way ANOVA, Bonferroni-correction | p = 0.1396 | ns | n = 8 |
| 0.00:Geo vs. 0.25:mCherry | Two-way ANOVA, Bonferroni-correction | p > 0.9999 | ns | n = 8 |
| 0.00:Geo vs. 0.25:HHF | Two-way ANOVA, Bonferroni-correction | p > 0.9999 | ns | n = 8 |
| 0.00:Geo vs. 0.25:Mx | Two-way ANOVA, Bonferroni-correction | p > 0.9999 | ns | n = 8 |
| 0.00:Geo vs. 0.25:Geo | Two-way ANOVA, Bonferroni-correction | p > 0.9999 | ns | n = 8 |
| 0.00:Geo vs. 0.50:mCherry | Two-way ANOVA, Bonferroni-correction | p > 0.9999 | ns | n = 8 |
| 0.00:Geo vs. 0.50:HHF | Two-way ANOVA, Bonferroni-correction | p > 0.9999 | ns | n = 8 |
| 0.00:Geo vs. 0.50:Mx | Two-way ANOVA, Bonferroni-correction | p > 0.9999 | ns | n = 8 |
| 0.00:Geo vs. 0.50:Geo | Two-way ANOVA, Bonferroni-correction | p > 0.9999 | ns | n = 8 |
| 0.25:mCherry vs. 0.25:HHF | Two-way ANOVA, Bonferroni-correction | p > 0.9999 | ns | n = 8 |

| | | | | |
|---|---|---|---|---|
| 0.25:mCherry vs. 0.25:Mx | Two-way ANOVA, Bonferroni-correction | p > 0.9999 | ns | n = 8 |
| 0.25:mCherry vs. 0.25:Geo | Two-way ANOVA, Bonferroni-correction | p > 0.9999 | ns | n = 8 |
| 0.25:mCherry vs. 0.50:mCherry | Two-way ANOVA, Bonferroni-correction | p > 0.9999 | ns | n = 8 |
| 0.25:mCherry vs. 0.50:HHF | Two-way ANOVA, Bonferroni-correction | p > 0.9999 | ns | n = 8 |
| 0.25:mCherry vs. 0.50:Mx | Two-way ANOVA, Bonferroni-correction | p > 0.9999 | ns | n = 8 |
| 0.25:mCherry vs. 0.50:Geo | Two-way ANOVA, Bonferroni-correction | p > 0.9999 | ns | n = 8 |
| 0.25:HHF vs. 0.25:Mx | Two-way ANOVA, Bonferroni-correction | p > 0.9999 | ns | n = 8 |
| 0.25:HHF vs. 0.25:Geo | Two-way ANOVA, Bonferroni-correction | p > 0.9999 | ns | n = 8 |
| 0.25:HHF vs. 0.50:mCherry | Two-way ANOVA, Bonferroni-correction | p > 0.9999 | ns | n = 8 |
| 0.25:HHF vs. 0.50:HHF | Two-way ANOVA, Bonferroni-correction | p > 0.9999 | ns | n = 8 |
| 0.25:HHF vs. 0.50:Mx | Two-way ANOVA, Bonferroni-correction | p > 0.9999 | ns | n = 8 |
| 0.25:HHF vs. 0.50:Geo | Two-way ANOVA, Bonferroni-correction | p > 0.9999 | ns | n = 8 |
| 0.25:Mx vs. 0.25:Geo | Two-way ANOVA, Bonferroni-correction | p > 0.9999 | ns | n = 8 |
| 0.25:Mx vs. 0.50:mCherry | Two-way ANOVA, Bonferroni-correction | p > 0.9999 | ns | n = 8 |
| 0.25:Mx vs. 0.50:HHF | Two-way ANOVA, Bonferroni-correction | p > 0.9999 | ns | n = 8 |
| 0.25:Mx vs. 0.50:Mx | Two-way ANOVA, Bonferroni-correction | p > 0.9999 | ns | n = 8 |
| 0.25:Mx vs. 0.50:Geo | Two-way ANOVA, Bonferroni-correction | p > 0.9999 | ns | n = 8 |
| 0.25:Geo vs. 0.50:mCherry | Two-way ANOVA, Bonferroni-correction | p > 0.9999 | ns | n = 8 |
| 0.25:Geo vs. 0.50:HHF | Two-way ANOVA, Bonferroni-correction | p > 0.9999 | ns | n = 8 |
| 0.25:Geo vs. 0.50:Mx | Two-way ANOVA, Bonferroni-correction | p > 0.9999 | ns | n = 8 |
| 0.25:Geo vs. 0.50:Geo | Two-way ANOVA, Bonferroni-correction | p > 0.9999 | ns | n = 8 |
| 0.50:mCherry vs. 0.50:HHF | Two-way ANOVA, Bonferroni-correction | p > 0.9999 | ns | n = 8 |

| | | | | |
|---|---|---|---|---|
| 0.50:mCherry vs. 0.50:Mx | Two-way ANOVA, Bonferroni-correction | p > 0.9999 | ns | n = 8 |
| 0.50:mCherry vs. 0.50:Geo | Two-way ANOVA, Bonferroni-correction | p > 0.9999 | ns | n = 8 |
| 0.50:HHF vs. 0.50:Mx | Two-way ANOVA, Bonferroni-correction | p > 0.9999 | ns | n = 8 |
| 0.50:HHF vs. 0.50:Geo | Two-way ANOVA, Bonferroni-correction | p > 0.9999 | ns | n = 8 |
| 0.50:Mx vs. 0.50:Geo | Two-way ANOVA, Bonferroni-correction | p > 0.9999 | ns | n = 8 |

### SUPPLEMENTARY REFERENCES


1. McHugh, C. A. *et al.* A virus capsid-like nanocompartment that stores iron and protects bacteria from oxidative stress. *EMBO J.* **33**, 1896–1911 (2014).
2. Giessen, T. W. & Silver, P. A. Widespread distribution of encapsulin nanocompartments reveals functional diversity. *Nat Microbiol* **2**, 17029 (2017).
3. Sigmund, F. *et al.* Bacterial encapsulins as orthogonal compartments for mammalian cell engineering. *Nat. Commun.* **9**, 1990 (2018).
